# Phenotypic analysis of an MLL-AF4 gene regulatory network reveals indirect CASP9 repression as a mode of inducing apoptosis resistance

**DOI:** 10.1101/2020.06.30.179796

**Authors:** Joe R. Harman, Ross Thorne, Max Jamilly, Marta Tapia, Nicholas T. Crump, Siobhan Rice, Ryan Beveridge, Edward Morrissey, Marella F.T.R de Bruijn, Irene Roberts, Anindita Roy, Tudor A. Fulga, Thomas A. Milne

**Affiliations:** MRC Molecular Haematology Unit, MRC Weatherall Institute of Molecular Medicine, Radcliffe Department of Medicine, University of Oxford, Oxford, OX3 9DS, UK; MRC Weatherall Institute of Molecular Medicine, Radcliffe Department of Medicine, University of Oxford, Oxford, OX3 9DS, UK; Marta Tapia, The Finsen Laboratory, Rigshospitalet, Faculty of Health Sciences, and Biotech Research and Innovation Centre (BRIC), Faculty of Health Sciences, University of Copenhagen, Copenhagen, Denmark; Department of Paediatrics, University of Oxford, Oxford, UK; Virus Screening Facility, MRC Weatherall Institute of Molecular Medicine, John Radcliffe Hospital, University of Oxford, Oxford, OX3 9DS, UK; Center for computational biology, Weatherall Institute of Molecular Medicine, University of Oxford, John Radcliffe Hospital, Oxford OX3 9DS, UK; NIHR Oxford Biomedical Research Centre Haematology Theme

## Abstract

Regulatory interactions mediated by transcription factors (TFs) make up complex networks that control cellular behavior. Fully understanding these gene regulatory networks (GRNs) offers greater insight into the consequences of disease-causing perturbations than studying single TF binding events in isolation. Chromosomal translocations of the *Mixed Lineage Leukemia gene* (*MLL*) produce MLL fusion proteins such as MLL-AF4, causing poor prognosis acute lymphoblastic leukemias (ALLs). MLL-AF4 is thought to drive leukemogenesis by directly binding to genes and inducing aberrant overexpression of key gene targets, including anti-apoptotic factors such as BCL-2. However, this model minimizes the potential for circuit generated regulatory outputs, including gene repression. To better understand the MLL-AF4 driven regulatory landscape, we integrated ChIP-seq, patient RNA-seq and CRISPR essentiality screens to generate a model GRN. This GRN identified several key transcription factors, including RUNX1, that regulate target genes using feed-forward loop and cascade motifs. We used CRISPR screening in the presence of the BCL-2 inhibitor venetoclax to identify functional impacts on apoptosis. This identified an MLL-AF4:RUNX1 cascade that represses *CASP9,* perturbation of which disrupts venetoclax induced apoptosis. This illustrates how our GRN can be used to better understand potential mechanisms of drug resistance acquisition.

**Graphical abstract caption:** A network model of the MLL-AF4 regulatory landscape identifies feed-forward loop and cascade motifs. Functional screening using CRISPR and venetoclax identified an MLL-AF4:RUNX1:*CASP9* repressive cascade that impairs drug-induced cell death.

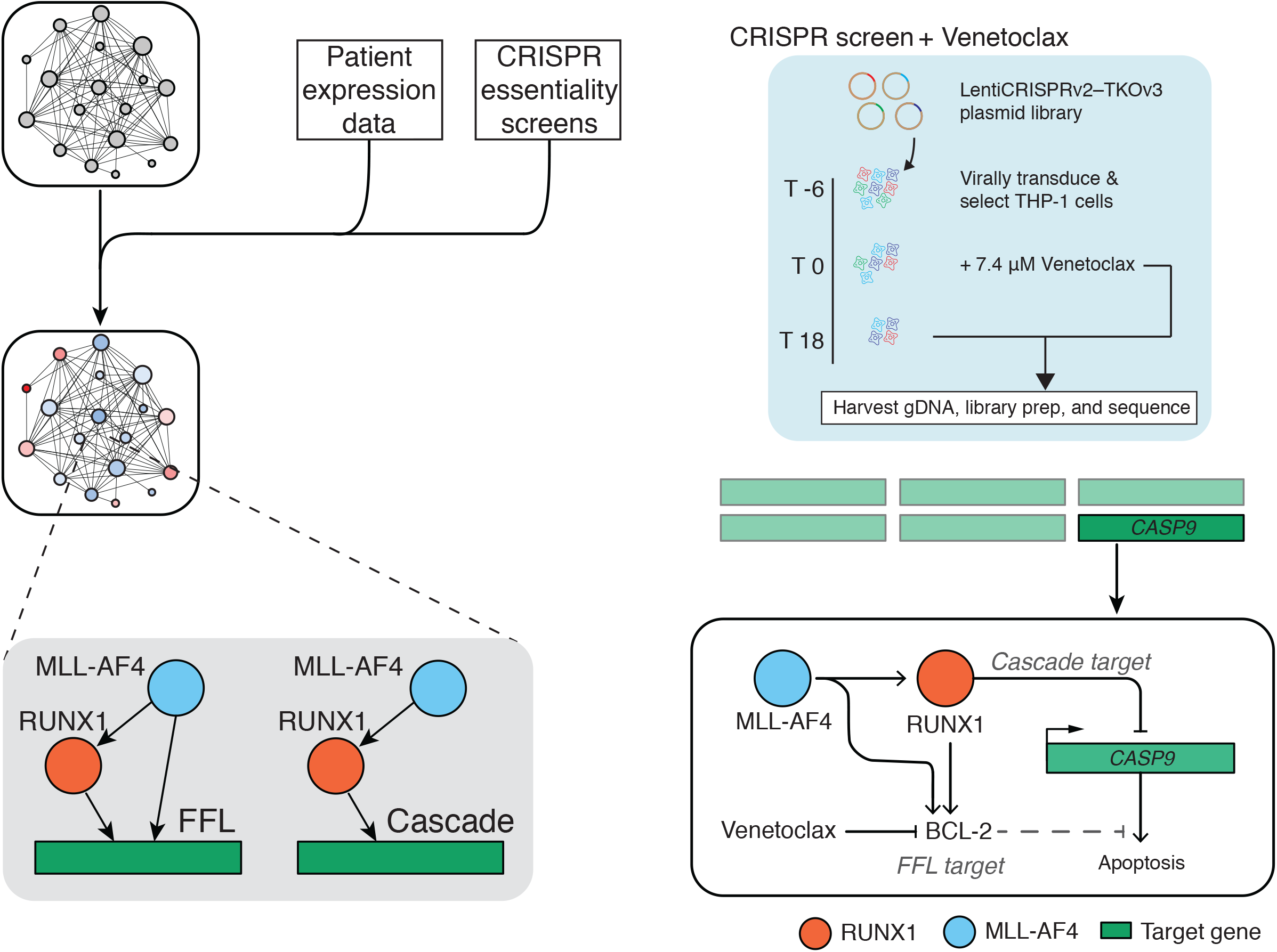

## INTRODUCTION

The regulated transcription of genes in eukaryotes is a core aspect of cellular behavior. Although there are often key individual genes that have to be appropriately regulated for normal tissue development, the coordinated regulation of entire sets of genes is required for the normal development of tissues, as well as for maintaining cellular homeostasis (1, 2). Disruption of these normal gene regulatory patterns can cause abnormal development and lead to human diseases such as leukemia (3). Gene regulatory patterns are controlled by the combinatorial activity of key sets of particular master transcription factors (TFs) (4, 5). A better model of this combinatorial code is necessary not only for understanding normal gene regulation, but also for understanding how these complex interactions are disrupted in human disease.

In order to better understand the combinatorial code of gene regulation driven by TFs, researchers have attempted to construct gene regulatory networks (GRNs) in the context of normal and abnormal biological systems. Many GRNs have been developed in single cell organisms, such as *Saccharomyces cerevisiae* and *Escherichia coli* (6–8), which formed the foundation for work from the Alon lab that made observations on the distinct structure of biological networks (8, 9). Following this work there have been many papers focusing on the consequence of network structure on interaction dynamics (10–13), or on the evolution of specific structural motifs (14, 15). GRNs have, in recent years, been applied to describe developmental systems such as neural crest development (16), hematopoietic specification (17) and T-lymphocyte specification (18). They have also been used to describe various cancers, including breast (19), prostate (20), colon (21), lymphoma (22) and leukemia (23).

Past attempts to understand network structure have focused on the overall connectivity of the system or on breaking down complex patterns into simple repeating units. Early development of network attempted to draw general conclusions about the structure of networks through the measure of centrality, which describes the relative number of connections of a node to the local network (24–26). Two such measures include degree centrality (the sum of connections to and from a node) and stress centrality (mapping the shortest paths between pairs of nodes in the network and quantifying how frequently hub nodes are involved in these shortest paths). Work from the Alon lab has suggested that network structure in biological systems can be reduced to repeating patterns of simple three-node motifs (9, 13). Particular patterns are enriched in GRNs compared to other types of networks. For example, one of the most prominent motifs is known as the feed-forward loop (FFL) (9, 13). Another recurring pattern is the TF cascade (or regulator chain), in which intermediate TFs propagate signal to indirect targets (7, 27). As compelling as these simple repeating motifs are, recognition of these patterns is a poor indicator of biological function in the absence of detailed functional experimentation (28) that cannot reasonably be performed genome wide. Thus, it has been difficult to use these simple patterns to make predictions or to better understand the connections that make up disease causing networks. In addition, GRNs can never be a complete representation of transcriptomic regulation. For example, while we present networks as a static representation of biology there is an underappreciated role of chaos in regulatory circuits (28), and there are questions regarding the adaptability of regulatory interactions to different environments (29), which cannot be truly modelled without extensive experimental testing. For these reasons, we suggest there is no one GRN applicable to all scenarios, and that they are better considered as a collection of possible pathways that describe TF interactions. We can then use these pathways to make predictions and build hypotheses that relate TF function to cell behaviors.

Leukemias driven by rearrangements of the *Mixed Lineage Leukemia* (*MLL*, also known as *KMT2A*) gene represent a very good example of a human disease caused by a single mutation that leads to dysregulated transcriptional networks. The most common MLL rearrangements (MLLr) are chromosome translocations that fuse MLL in frame to one of a wide number of partner genes, creating novel fusion proteins (MLL-FPs) (30). Generally, MLL-FP driven leukemias do not respond well to treatment and have a very poor prognosis (31, 32). MLL-FPs drive both acute myeloid leukemia (AML) and acute lymphoblastic leukemia (ALL), and are known to switch lineage. MLLr ALL can relapse after treatment as an AML that derives from the original leukemic clone (33, 34), suggesting that a single MLL-FP is able to drive both ALL and AML transcriptional networks. MLL-FPs that arise from these translocations bind to target genes and promote transcription through the recruitment of a large transcription elongation complex (35). One key part of MLL-FP function involves recruiting the Disruptor of Telomeric Silencing 1-like (DOT1L) protein and increasing histone H3 lysine 79 methylation (H3K79me2/3) levels (36–45). Targets of these MLL-FPs include key TF genes such as *HOXA9* and *RUNX1* (40, 46), perturbation of which may lead to further transcriptomic disruption. One of the most common MLL rearrangements is a t(4;11) (q21;q23) chromosomal translocation, fusing *KMT2A* with *AF4* (also known as *MLLT2*), to produce an MLL-AF4 fusion protein, most commonly associated with ALL (30). MLL-AF4 leukemias have few cooperating mutations (47–49) which suggests this chromosomal translocation by itself is sufficient for leukemic transformation, and thus MLL-AF4 leukemias represent an ideal system for understanding how a single perturbation can deregulate a transcriptional network in human disease.

The general approach in the field towards transcriptomic profiling of leukemias has been to consider differentially expressed genes (DEGs) that are not directly bound by MLL-AF4 to reflect either indirect downstream effects or transcriptional noise caused by changes in cellular context, and therefore not directly related to the function of the MLL-FP. However, since MLL-AF4 and other MLL-FPs directly regulate the expression of key TFs, this ignores the wider transcriptional network regulated by MLL-AF4, with downstream TFs potentially connecting these “indirect” targets to direct MLL-FP activity (Figure 1A). Here we used systems approaches to describe the larger transcriptional network driven by MLL-AF4 that facilitates indirect control of downstream targets. We then dissected these complex interactions to identify network motifs that could be directly tested experimentally. Our approach was to use GRNs as a way of organizing complex data into simple graph structures to create testable hypotheses about key downstream targets that can be used to help interpret patient data or functional screens.

**Figure 1.**
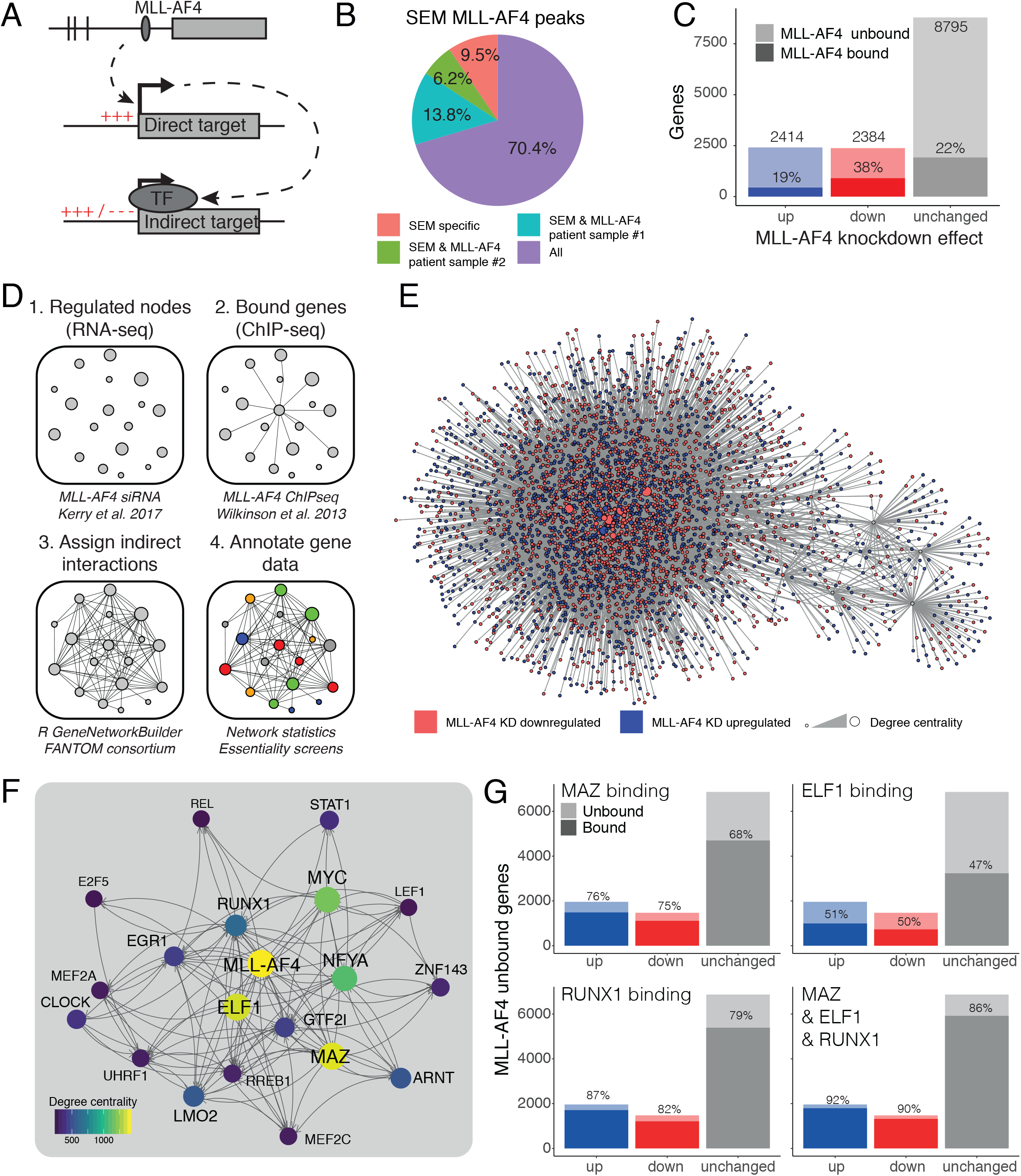
Developing a GRN model to assess indirect regulatory effects of MLL-AF4. (**A**) Schematic illustrating concept of MLL-AF4 targeting of TF genes, leading to subsequent TF protein expression and downstream regulation of indirect targets. (**B**) Pie chart of MLL-AF4 target genes in SEM cells (defined by nearest annotated promoter in ChIP-seq), describing overlap with MLL-AF4 ChIP-seq target genes from patient samples. (**C**) DEGs from nascent RNA-seq after 96 hours’ MLL-AF4 siRNA knockdown. Differential expression is defined as FDR < 0.05. (n=3). Shaded area represents genes bound by MLL-AF4 ChIP-seq. (**D**) Illustration of workflow used to generate a GRN from nascent RNA-seq and ChIP-seq data. (**E**) Visualization of whole network. Nodes in red represent genes downregulated upon MLL-AF4 KD, while blue represents upregulated genes. Circle size represents degree centrality. (**F**) Top 20 genes of the MLL-AF4 GRN by degree centrality. Lines (edges) indicate predicted interaction from protein (source) to gene locus (target), with arrowheads pointing towards downstream nodes. (**G**) DEGs after MLL-AF4 knockdown that are unbound by MLL-AF4, as highlighted in (**C**). Shaded areas represent proportion of genes bound by MAZ, ELF1 or RUNX1 ChIP-seq, as indicated.

One example of a known key pathway in MLL leukemias is the regulation of apoptosis through the BCL-2 family of proteins. Recent work on MLL-FP leukemias has shown that BCL-2 protein levels are abnormally high in MLL-AF4 ALL, due to direct regulation of the *BCL2* gene by MLL-AF4 (50). The BCL-2 family is important for controlling the intrinsic apoptotic pathway by integrating information from both extracellular and intracellular signaling pathways. This is achieved through the delicate balance between directly interacting pro- and anti-apoptotic BCL-2 family proteins (51, 52). High levels of proteins such as BCL-2, BCL-XL and MCL-1 block apoptosis. The expression of BCL-2 family proteins is perturbed in multiple types of cancers, including leukemias, and is associated with disease progression and resistance to chemotherapy (53, 54). Because of this, much work has been put into developing inhibitors such as venetoclax (ABT-199) that specifically target the BCL-2 protein itself. Interestingly, MLL-FP leukemias are particularly sensitive to treatment with venetoclax (50, 55, 56). However, due to the complexity of the BCL-2 family pathway, there are many ways in which cells are able to develop resistance to inhibitors such as venetoclax. In addition, there is the potential for an MLL-AF4 driven GRN to control regulation of the BCL-2 pathway in ways that are not easy to directly predict from protein:DNA binding and expression patterns alone. As direct MLL-AF4 binding activity is not sufficient to determine the regulatory scope of the fusion protein, our goal was to establish whether simple MLL-AF4 GRN motifs could be used to better understand drug resistance pathways in MLL-FP leukemia cells, such as the BCL-2 apoptotic pathway.

Using SEM cells (an MLL-AF4 ALL cell line) we created a GRN using RNA-seq, ChIP-seq, and TF interaction data from the FANTOM consortium (57). Our model shows that just three highly central TFs, RUNX1, ELF1 and MAZ, are capable of accounting for almost all of the indirect MLL-AF4 regulation and are ubiquitously expressed across AML and ALL patient RNA-seq samples. *RUNX1*, a known target of MLL-AF4 (46), is essential for the viability of MLL-FP cell lines in CRISPR screens (58). Using a phenotypic approach, we looked for MLL-AF4:RUNX1 GRN motifs involved in apoptosis and found, among others, an MLL-AF4:RUNX1-mediated cascade repressing *CASP9*, which encodes the pro-apoptotic effector protein Caspase-9. Using venetoclax to drive apoptosis in combination with a CRISPR knockout screen in the THP-1 cell line (MLL-AF9 AML) revealed several genes within the GRN that reduced or exacerbated drug-induced apoptosis in these cells. Amongst these genes, we found that CRISPR targeting of *CASP9* promotes survival. We confirm that MLL-AF4 does not appear to repress *CASP9* directly but does so via intermediate TFs such as RUNX1. This highlights the capacity for MLL-AF4 to alter cell behaviors such as apoptosis not just through direct regulation of gene targets, but also by perturbation of the wider transcriptional network. We propose that this MLL-AF4:RUNX1:CASP9 pathway, among others identified in our network, highlights key regulatory circuits in leukemia cells and provides possible pathways by which MLL-FP leukemias may escape therapeutic intervention.

## MATERIAL AND METHODS

### Cell line culture

SEM cells, an MLL-AF4 B cell ALL line (59), were purchased from DSMZ (www.cell-lines.de). SEM cells were cultured in Iscove’s modified Dulbecco’s medium (IMDM) with 10% fetal bovine serum (FBS) and 1x GlutaMAX, with cell density maintained between 5×10^5^/ml and 2×10^6^/ml. THP-1 cells, an MLL-AF9 AML cell line, were purchased from ATCC (www.lgcstandards-atcc.org). THP-1 cells were cultured in RPMI-1640 with 10% FBS and 1x GlutaMAX, with cell density maintained between 5×10^5^/ml and 1.5×10^6^/ml. Cells were confirmed to be free of mycoplasma.

### Patient samples

MLL-AF4 patient sample #1 is described in Kerry et al. 2017 (60). MLL-AF4 patient sample #2 is a primary diagnostic bone marrow sample from a 6-year-old child with ALL, obtained from the Bloodwise Childhood Leukaemia Cell Bank, UK (REC: 16/SW/0219). Samples were anonymized at source, assigned a unique study number and linked.

### siRNA knockdowns

1×10^7^ SEM cells in log phase growth were transfected with 10 μl 20 μM siRNA by electroporation. Cells were then seeded into medium at 1×10^6^/ml. For western blot and qRT-PCR analysis cells were re-transfected as above 24 hours later (RUNX1 siRNA) or 48 hours later (MLL-AF4 siRNA, MAZ siRNA). For RNA-seq analysis, SEM cells were transfected with RUNX1 siRNA for 96 hours. The following siRNA were used: MLL-AF4 siRNA (siMA6) and scrambled control (siMM) (61); RUNX1 (Dharmacon ON-TARGETplus single siRNA, J-003926-05); MAZ (Ambion Silencer Select, s8543); Non-targeting controls (Dharmacon ON-TARGETplus non-targeting pool, D-001810-10-20).

### Western blots

Proteins for western blot analysis were extracted using BC300 lysis buffer with protease inhibitors. Samples were loaded onto a 4-12% bis-tris gel and blotted using polyvinylidene fluoride membrane for 1 hour at 100 V using a tris-glycine transfer buffer. Membranes were incubated with primary antibody in 5% milk/TBS-T (5% BSA/TBS-T for anti-RUNX1) overnight at 4°C and subsequently incubated with anti-Rabbit IgG peroxidase antibody (1/10,000, Sigma A6667) for 1 hour at room temperature. Primary antibodies were all raised in rabbit and include anti-RUNX1 (1/5000, Cell Signaling 4334S); anti-Caspase-9 (1/5000, Abcam ab202068); anti-MAZ (1/5000, Bethyl A301-652A); anti-GAPDH (1/10,000, Bethyl A300-641A); anti-Vinculin (1/50,000, Abcam ab129002).

### qRT-PCR

Total RNA was extracted using the RNeasy Mini kit (Qiagen) 24 or 48 hours post siRNA knockdown. cDNA was generated using SuperScript III Reverse Transcriptase (Life Technologies) with random hexamer primers. qRT-PCR analysis was performed using TaqMan or SYBR probes and analyzed with the ΔΔCt method normalizing to the housekeeping gene *GAPDH* (Hs03929097_g1). TaqMan probes were used for *RUNX1* (Hs00231079_m1); *CASP9* (Hs00609647_m1), *BCL2* (Hs00608023_m1), *CDK6* (Hs01026371_m1), *PROM1* (Hs01009257_m1), *MYC* (Hs0015348_m1), *ELF1* (Hs00608023_m1), *GNAQ* (Hs01586104_m). SYBR probes were used for *MAZ* and *MLL-AF4* (For primer sequences see Supplementary Table S1). Statistical analyses were performed using two-sided student’s t-test.

### Nascent RNA-seq library prep

Nascent RNA-seq was performed 96 hours post *RUNX1* knockdown. Details on nascent RNA protocol is as previously published (60). Briefly, 1×10^8^ SEM cells were treated with 500 μM 4-thiouridine (4-SU) for 1 hour. Cells were lysed with Trizol (Life Technologies) and total RNA was precipitated and DNase I-treated. 4-SU-incorporated RNA was purified by biotinylation and streptavidin bead pulldown. DNA libraries were prepared from nascent RNA using the NEB Next Ultra Directional RNA library preparation kit. Samples were sequenced by paired-end sequencing using a NextSeq 500 (Illumina).

### RNA-seq analysis

FASTQ files were quality checked using FastQC (v0.11.4). Adapters and poor-quality bases were then trimmed from the reads using trim_galore (v0.4.1). Paired end reads were mapped to the human genome assembly (hg19) using the STAR aligner (v2.4.2). PCR duplicates were removed using picard-tools MarkDuplicates (v1.83). Mapped reads were then quantified over gene exons using subread featureCounts (v1.6.2) to measure gene expression levels. Statistical analysis was performed in R using the edgeR package (62), and P values were corrected using the Benjamini & Hochberg method to acquire False Discovery Rates (FDR). Genes were considered differentially expressed if they had an FDR of less than 0.05. Differentially expressed genes were analyzed for enriched GO terms and reactome pathways using PANTHER (v. 15) (63). Processed expression tables were used for published patient RNA-seq analyses (64–66), with GEO accession numbers reported in Supplementary Table S2. RNA-seq expression Pearson correlations were calculated using R.

### ChIP-seq assay

1×10^8^ cells were fixed and sonicated using a Covaris (Woburn, MA, USA) according to the manufacturer’s recommendations. A mixture of magnetic Protein A and Protein G Dynabeads (Life Technologies) were used to isolate Ab:chromatin complexes, which were washed three times with a solution of 50 mM HEPES-KOH, pH 7.6, 500 mM LiCl, 1 mM EDTA, 1% NP-40, and 0.7% Na deoxycholate. Antibodies used include anti-MAZ (Bethyl, A301-652A), anti-MLL-N (Bethyl, A300-086A) and anti-AF4-C (Abcam ab31812). The samples were washed with Tris-EDTA, eluted, and treated with RNase A and proteinase K. DNA was then purified with a Qiagen PCR purification kit. Libraries were generated using NEBnext Ultra DNA library preparation kit for Illumina (NEB). Libraries were sequenced by paired-end sequencing using a NextSeq 500 (Illumina).

### ChIP-seq analysis

To analyze ChIP-seq data quality control of FASTQ reads, alignment, PCR duplicate filtering, blacklisted region filtering, and UCSC data hub generation was performed using an in-house pipeline (https://github.com/Hughes-Genome-Group/NGseqBasic/releases). Briefly, the quality of FASTQ files were checked using FastQC (v0.11.4) and then mapped using Bowtie (v1.0.0) against the human genome assembly (hg19). Unmapped reads were trimmed with trim_galore (v0.3.1) and mapped again. Short unmapped reads from this step were combined using flash (v1.2.8) and mapped again. PCR duplicates were removed using samtools rmdup (v0.1.19). Any reads mapping to Duke blacklisted regions (UCSC) were removed using bedtools (v2.17.0). Directories of ChIP-seq tags (reads) were generated from the SAM files using Homer makeTagDirectory (v4.7). To visualize on UCSC the makeBigWig.pl command was used to generate bigwig files normalizing tag counts to tags per 10 million tags. MLL-N and AF4-C peaks were called using SeqMonk (v0.24.1) and RUNX1, ELF1 and MAZ peaks were called using Homer findPeaks (-style factor), with both tools using an input track for background correction. MLL-AF4 binding was defined as overlapping MLL-N and AF4-C peaks. Peaks were associated with the nearest promoter using Homer annotatePeaks.pl (v4.8). Metagene profiles were generated with Homer annotatePeaks.pl and visualized using R. ChIP-seq heatmaps were generated using DeepTools (v3.0.1) computeMatrix centered on MLL-AF4 or RUNX1 peaks with a 6 kb window. RUNX1 motifs were identified genome wide using Homer scanMotifGenomeWide.pl (v4.8) with RUNX1 HPC7 motifs from the Homer motif database.

### GRN creation and analysis

GRNs were created on the basis of a central node: MLL-AF4, RUNX1, BRD4 or DOT1L. The GRN construction involved 4 steps. 1). To determine the regulatory scope of the central node we used nascent RNA-seq data obtained from SEM cells after 96 hours’ *MLL-AF4* (60) and 96 hours’ *RUNX1* siRNA knockdowns, 7 days’ EPZ-5676 treatment (a sufficient duration to produce near complete loss of H3K79me3) to inhibit DOT1L (38), and 1.5 hours’ IBET-151 treatment (a sufficient duration to observe immediate effects of BET domain inhibition) to inhibit BRD4 (67). 2). Direct interactions were established using ChIP-seq peaks corresponding to the central node which were annotated to the nearest gene promoter using Homer annotatePeaks.pl (v4.7). These annotations were assumed to represent direct protein-gene interactions. For RUNX1 we additionally filtered the ChIP-seq peaks for those that overlap SEM enhancers, identified as H3K27ac overlapped with H3K4me1 ChIP-seq peaks. 3). Indirect interactions in the GRNs were integrated with the R package GeneNetworkBuilder (v1.26.1) using TF interaction data from the FANTOM consortium (57). TF interactions in this underlying dataset that do not explain connectivity between the central node and its direct or indirect regulatory targets were excluded from the resulting GRN. 4). Overlaps between GRNs were tested using Fisher’s exact test. Nodes of the network were annotated using ALL and AML patient RNA-seq (64–66), CRISPR screens(58, 68), and measures of node centrality. Degree and stress centralities were calculated using the R packages igraph (v1.2.4.1) and sna (v2.4).

### Patient sub-network creation and analysis

We made use of published RNA-seq data, derived from leukemic blasts from patients and normal fetal bone marrow (FBM) and B cell populations, to create subgraphs of the MLL-AF4 GRN. We used published processed data from MLLr ALL patients (66), AML patients with a range of chromosomal abnormalities (65), and normal FBM (n=3 samples) hematopoietic stem and progenitor cell (HSPC) populations and B cells (64). Patient sub-networks were derived by filtering the *MLL-AF4* knockdown RNA-seq for genes expressed in each patient RNA-seq sample and reperforming the GRN creation workflow. Expressed genes are defined as those with greater than 1.5 log_2_ TPM (see Supplementary Figure S2A). Node presence and absence across all sub-networks were converted into a binary matrix, and UMAP dimensionality reduction was performed using this matrix to identify clusters of nodes using R.

### CRISPR screen generation

Pooled lentiviral CRISPR knockout screening was conducted with the TKOv3 genome-wide sgRNA library as described previously (69) with the following modifications. To generate the TKO v3 lentiviral library HEK293T cells were plated at 1.4 × 10^7^ cells per 15 cm dish (DMEM, 10% FCS, penicillin/streptomycin, L-glutamine), the following day they were transfected with the pooled library plasmid, pMD2.G and psPAX2 plasmids using PEI Pro (Polyplus Transfection). 6 hours post transfection the medium was changed. Harvests were collected 48- and 72-hours post transfection, combined and filtered through a 0.45 μM cellulose acetate filter. Filtered viral supernatants were aliquoted and stored at −80°C. The single-vector TKOv3-lentiCRISPRv2 library was a gift from Jason Moffat (Addgene #90294) (70). The screen was performed in duplicate each using fresh cells and lentivirus. Library representation was maintained at a minimum of 250 cells/sgRNA throughout the screen. THP-1 cells were transduced with TKOv3 lentivirus at MOI=0.3 to ensure single guide/cell transduction. After 72 hours, transduced cells were selected with 2 μg/ml puromycin. After 72 hours’ expansion, 1.8×10^7^ viable cells were harvested for NGS to serve as a baseline (day 0 of drug treatment). A further 4.7×10^8^ viable cells were cultured with 7.4 μM venetoclax (ABT-199, Stratech) for 18 days (six passages), with fresh venetoclax used at each passage. We used the Annexin-V/PI apoptosis assay to determine that 7.4 μM is the half-maximal cytotoxic concentration of venetoclax in THP-1 cells. On day 18, 1.8×10^7^ viable cells were harvested for NGS. Cells were centrifuged at 100g for 10 minutes to separate viable cells from dead cells. gDNA was isolated using the QIAamp DNA Blood Maxi Kit (Qiagen). sgRNA libraries were generated using two-step PCR and sequenced by paired-end sequencing using a NextSeq 500 (Illumina).

### CRISPR screen analysis

sgRNA forward and reverse NGS reads were merged using BBMerge with default quality filtering. >95% of reads were successfully merged. sgRNAs were aligned to the TKOv3 library using CRISPressoCount (71) with a quality score threshold of 30 to generate sgRNA counts for day 0 and day 18. This pipeline yielded a mean of ≥188 aligned reads/sgRNA. ≥99.6% of all sgRNAs were detected in every sample. sgRNA counts were analyzed with MAGeCK-RRA (v0.5.7) (72) using copy number variation (CNV) data for THP-1 cells downloaded from the Cancer Cell Line Encyclopedia as described previously (73). Robust rank aggregation was performed with 10,000 permutations.

### Annexin V/PI assay

Annexin V/PI assay was used to determine viability after venetoclax treatment, as described previously (74, 75). THP-1 cells were treated for 48 hours with a range of venetoclax concentrations. THP-1 cells were then harvested, washed in PBS, and incubated in 50 μl annexin-V binding buffer (Biolegend) containing FITC-conjugated annexin-V (Biolegend) and propidium iodide (PI, ChemoMetec) for 15 minutes on ice. Cells were diluted to a final volume of 150 μl and analyzed using an Attune NxT flow cytometer (Thermo Fisher).

### CellTiter-Glo

To validate CRISPR screen hits, individual sgRNA sequences were cloned into lentiCRISPRv2, a gift from Feng Zhang (Addgene #52961), and used to target SEM or THP-1 cells. CRISPR-Cas9 edited SEM or THP-1 cells were treated for 48 hours with 2.5 μM or 20 μM venetoclax, respectively. Cell viability of these cells were assayed using the CellTiter-Glo (CTG) luminescence assay. Cell suspensions were mixed at a 1:1 ratio with CTG substrate, and luminescence was detected using a BMG FLUOstar OPTIMA plate reader. Viability for THP-1 cells were represented relative to *AAVS1* targeting sgRNA (76). Statistical analysis of CTG luminescence was performed using two-sided student’s t-test.

## RESULTS

### The MLL-AF4 fusion protein controls a wider transcription factor network through targeting key transcription factors

We have previously characterized the genome wide binding of MLL-AF4 (60) in SEM cells, an MLL-AF4 cell line originally derived from a patient (59). We annotated MLL-AF4 ChIP-seq peaks with the nearest gene promoter, which we refer to as MLL-AF4 bound genes, to predict functional targets of the fusion protein. It has been observed that cell lines show significant changes in gene regulation, compared with primary tissues (77), which may in part be due to epigenetic changes and altered genome wide protein:DNA binding profiles. To validate how representative our MLL-AF4 ChIP-seq is of primary tissue behavior, we overlapped the MLL-AF4 bound genes in SEM cells with bound genes in two different MLL-AF4 ALL patient samples ((60) and see methods) (Figure 1B). The majority of SEM MLL-AF4 bound targets show overlap with both MLL-AF4 patient samples (70%), which is sufficient to conclude that the SEM data is representative of a core set of MLL-AF4 DNA binding interactions. Further, using an antibody against the N terminus of the MLL-AF4 fusion protein (MLL-N), comparing MLL-N ChIP-seq signal at promoters shows a Spearman’s rank correlation of 0.73 and 0.74 between SEM and each of the patient samples. To determine what transcriptional networks MLL-AF4 controls, we analyzed nascent RNA-seq following MLL-AF4 siRNA knockdown for 96 hours (60) to identify DEGs. Considering the direct role of MLL-AF4 complex components in transcriptional activation (35, 37, 42, 43, 60, 78, 79), we might expect that the RNA-seq analysis would show a bias towards downregulation of genes upon knockdown. However, we identified 2414 upregulated and 2384 downregulated genes, showing no bias either way (Figure 1C). MLL-AF4 is often considered to be the main driver of leukemogenesis, therefore we might also expect that the majority of observed gene expression changes would be in genes directly bound by MLL-AF4. However, overlapping the DEGs with SEM MLL-AF4 ChIP-seq (46) reveals that only 19% of upregulated genes (450 genes) and 38% of downregulated genes (908 genes) are bound by MLL-AF4. The majority of these DEGs may therefore be regulated not by MLL-AF4 itself, but instead by intermediate TFs whose expression is regulated by the fusion protein (Figure 1A).

To explore this possibility, we constructed an MLL-AF4 GRN to better understand the wider regulatory landscape of the fusion protein and the connectivity towards MLL-AF4-unbound targets. In brief, we took all DEGs upon knockdown of MLL-AF4 to be the regulatory scope of the fusion protein (Fig 1D, box 1) and assigned direct interactions using the MLL-AF4 ChIP-seq data (Fig 1D, box 2). We then used TF interaction data from the FANTOM consortium (57) to incorporate regulatory interactions that connect differentially expressed TFs to downstream targets (Fig 1D, box 3). Genes were annotated with additional information from published datasets (58, 64–66, 68) (Fig 1D, box 4) to assign functional importance to each node of the network (Figure 1D, E, Supplementary Data S1).

The top nodes of this GRN, when ranked by degree centrality (the number of connections to and from a gene) include several key TFs such as ELF1, MAZ and RUNX1 (Figure 1F). We have previously shown a potential role for ELF1 and RUNX1 in MLL-AF4 leukemias (38, 46). Using previously published ChIP-seq data for ELF1 and RUNX1 (38, 46), and ChIP-seq generated for MAZ in SEM cells, we found that direct binding of any of these factors has the potential to account for the majority of differential effects of the MLL-AF4 knockdown. Using all of these factors together has the capacity to account for 92% and 90% of up- and downregulated genes that are unbound by MLL-AF4 (Figure 1G). We also found that MLL-AF4 binds to just 18 genes that are not bound by these three TFs, while RUNX1 has the most unique annotations with 306 gene targets (Supplementary Figure S1A), suggesting that the complex interplay of multiple TFs may determine the expression profile of most genes within the GRN. While DNA binding does not necessarily imply strong regulation of bound targets, this observation provides a potential mechanism by which MLL-AF4 could regulate indirect targets.

MLL-AF4 functions by assembling a large transcription elongation complex at gene targets, which includes the H3K79 methyltransferase DOT1L (36, 38, 60, 80). To test our network, we therefore compared it to a network centered around DOT1L. We also compared this to a network constructed using BRD4, a protein which is commonly associated with gene activation in MLL leukemias (81, 82) but which is not a direct component of the MLL-AF4 complex (37, 43, 78). For the DOT1L model we used nascent RNA-seq of SEM cells treated for 7 days with the DOT1L inhibitor (DOT1Li) EPZ-5676 (38, 83), along with H3K79me3 ChIP-seq (38) as a proxy for DOT1L binding (Supplementary Figure S1B). The BRD4 network made use of published nascent RNA-seq of SEM cells treated with IBET-151 for 1.5 hours to inhibit BRD4 binding to chromatin (67) and BRD4 ChIP-seq (60) (Supplementary Figure S1C). We found that, while many of the MLL-AF4 GRN nodes are affected by IBET and DOT1Li treatment, not all of the most central nodes are impacted (Supplementary Figure S1D). Further, overlapping the MLL-AF4 network with the DOT1L and BRD4 GRNs shows a greater number of overlapping nodes with BRD4 than DOT1L networks (2576 and 1363 respectively) (Supplementary Figure S1E). However, the MLL-AF4 GRN overlaps with a greater proportion of the DOT1L network than BRD4 (47.7% and 31.5% respectively), which is reflected in a greater statistical significance in overlap and indicates greater specificity of DOT1Li for the MLL-AF4 regulatory landscape (Supplementary Figure S1F). This matches our expectation that a core complex component should have a closer association with MLL-AF4 behavior.

### ALL and AML patient sub-networks highlight core, central transcription factors

To test the applicability of our model to leukemia in patients we set out to integrate patient RNA-seq datasets. We used published data from MLLr ALL patients (66), AML patients with a range of chromosomal abnormalities (65), and hematopoietic stem and progenitor cell (HSPC) populations and B cells from normal human fetal bone marrow (FBM; n=3 samples) (64). Our integration approach was to generate individual patient-specific sub-networks derived from our SEM MLL-AF4 GRN, by filtering the SEM MLL-AF4 RNA-seq results by genes expressed within each patient sample, and then re-processing the GRN workflow (Figure 2A). The level of expression at which a gene was considered active was 1.5 log_2_ TPM (Supplementary Figure S2A). The patient sub-networks were then collated and summarized as a binary matrix, describing node presence or absence across each patient, in the case of the ALL and AML datasets, and each individual sample in the case of the FBM dataset. Using this matrix, we calculated the frequency at which a node is expressed in each ALL, AML and FBM datasets and found that the most central nodes of the SEM MLL-AF4 GRN (degree centrality > 500, 8/3850 nodes) are constitutively active in both the ALL and AML datasets (Figure 2B). Strikingly, the most central nodes are also bound by MLL-AF4 in two ALL patient samples (see Figure 1B), indicating that they are commonly upregulated by MLL-AF4 in leukemia (Supplementary Figure S2B). The consistent expression of these highly connected TFs points towards a core role, not only in the SEM MLL-AF4 transcriptional network, but in all ALL and AML leukemias. We noted that the level of node conservation was reduced in the normal FBM samples compared to the leukemia dataset, particularly for the more central nodes (Figure 2B). For example, MAZ, which is expressed in all ALL and AML samples, is completely silent in the FBM dataset. This is consistent with the concept that the GRN is driven by MLL-AF4 to increase, or enable constitutive activation of, TF circuits. Therefore, these circuits represent common regulatory wiring that enables an oncogenic transcription program.

**Figure 2.**
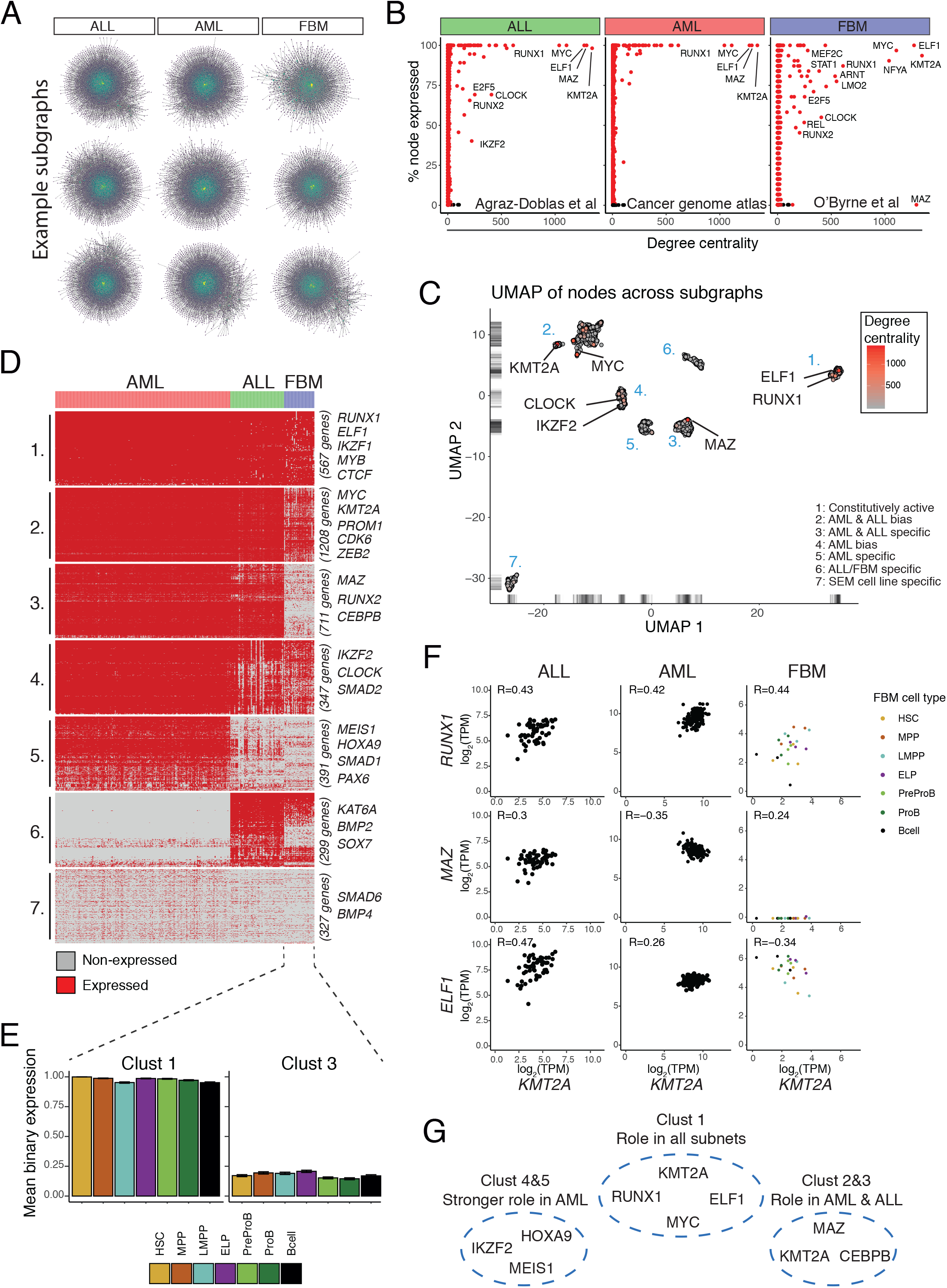
Patient sub-network analysis highlights core transcription factors. (**A**) Example sub-networks generated by filtering DEGs following MLL-AF4 KD in SEM cells by expressed genes in patient RNA-seq datasets (64–66), and running through the same pipeline as outlined in Figure 1D. For ALL and AML datasets each sub-network represents an individual patient. For FBM each sub-network represents an individual sample and cell type. Expressed genes are defined as log_2_ TPM > 1.5. Subgraphs and patient data were then summarized as a binary matrix describing whether each GRN node is expressed in each sample. (**B**) Using the binary expression in (**A**), the percentage of samples in each RNA-seq dataset that express each gene is plotted against degree centrality of the node in the MLL-AF4 GRN. (**C**) UMAP dimensionality reduction of the binary matrix created from (**A**). Genes are colored based on their degree centrality within the SEM GRN. (**D**) Heatmap of the binary expression matrix from (**A**), separated into seven clusters based on the gene groups seen in (**C**). Rows represent individual genes; columns represent individual samples. Red indicates gene expression while grey indicates no expression. (**E**) Scatter plots of select genes, plotted from log_2_(TPM) expression data from the ALL, AML and FBM datasets. (**F**) Box plots of FBM cell populations showing mean binary expression in clusters 1 and 3. (**G**) A summary of conclusions regarding key gene clusters in this analysis.

To further explore the primary RNA-seq material, we used the dimensionality reduction method UMAP to compare expression of each GRN node across the samples. This identified seven clusters of genes (Figure 2C-D, Supplementary Data S2). Cluster 1 shows active nodes across all samples, and includes several highly central nodes, such as *RUNX1* and *ELF1.* These genes are enriched for hematopoietic and differentiation processes (Supplementary Figure S2D). Cluster 2 instead shows slightly decreased node representation in the FBM dataset, which includes other highly central nodes including *KMT2A* and *MYC*. Note that *KMT2A* here could represent expression from normal *KMT2A*, but in the case of MLLr leukemias will also include MLL-AF4 expression. Interestingly, the *PROM1* gene (coding for CD133) is in cluster 2, which matches previous work identifying *PROM1* as an overexpressed target of MLL-AF4 that is also more generally expressed in cancer stem cells (84–86). Cluster 3 contains nodes that are specific to AML and ALL datasets, with almost no expression in FBM. As expected from Figure 2B*, MAZ* is in this cluster. MAZ has recently been implicated as having a role in the transcriptional network controlling embryonic hematopoietic stem cell (HSC) ontogeny, and from our data appears to be inactive in FBM. The reactivation of MAZ in leukemia may imply that MLL-AF4 co-opts some embryonic specific regulatory systems (87). To confirm that cluster 3 was not masking expression differences between the multiple cell types included in the FBM data we plotted the mean binary expression within each cell type, demonstrating similar expression frequencies across cell types (Figure 2E). Cluster 4 is biased towards nodes expressed in the AML dataset, and shows active expression of *IKZF2, CLOCK* and *CEBPB,* while cluster 5 is AML-specific and contains *MEIS1 and HOXA9* (Supplementary Figure S2C). Cluster 5 is also enriched for cell adhesion and chromatin organization factors. It is interesting that an AML-specific cluster was identified, considering the base network is generated using SEM cells, an ALL line, though this is a minority of the total GRN nodes (19%). In contrast, cluster 6 shows a bias towards ALL, and contains genes associated with response to external stimulus and protein phosphorylation (Supplementary Figure S2D). This cluster also contains a mix of ALL- and FBM-biased genes. Those specific to ALL include *SOX7*, *CD70*, *LAMP5* and *KAT6A*, while those shared between ALL and FBM include *HIST1H2AH* and *KAT7*. Cluster 7 contains few or no nodes expressed in any of the datasets, and likely represents nodes which are unique to the SEM network and therefore are not clinically relevant. This cluster is enriched for terms related to cell cycle and stress response, and likely represents a regulatory program specific to immortalized cell lines (Supplementary Figure S2D).

Three of the most central nodes in the network, *RUNX1*, *ELF1* and *MAZ*, are frequently expressed in ALL and AML datasets (Figure 2B). *RUNX1* correlates with *KMT2A* across all datasets, while there are weaker correlations for *MAZ* and *ELF1* (Figure 2F). Interestingly, *ELF1* appears to show stronger correlation with *KMT2A* in the ALL dataset over AML, and it appears to be inversely correlated with *KMT2A* in the *FBM* data. We also see clearly that the lack of *MAZ* nodes in the FBM subnetworks is not due to our definition of expressed genes (Supplementary Figure S2A), as samples show few to no transcripts (Figure 2F). Using these data we were able to group nodes of the MLL-AF4 GRN into different modules that describe their relationship with patient data, revealing genes representing an ubiquitous circuit present across all samples, those with a general bias towards leukemic samples over normal FBM, and those with a more specific role in AML patients (Figure 2G).

### Core GRN factors are correlated with MLL-AF4 binding

The highly central TFs that we have identified may have a distinct or cooperative role with MLL-AF4. Therefore, to explore the interplay of these factors with MLL-AF4 we performed a more detailed analysis of genome-wide ChIP-seq binding profiles. MLL-AF4 binds to gene promoters, and at a subset of targets spreads into the gene body. These spreading targets are sensitive to DOT1L inhibition (60). At the *GNAQ* locus, the MLL-AF4 spreading peak is enriched for RUNX1, ELF1 and MAZ ChIP-seq signal, with multiple peaks observed (Figure 3A). To quantify this more generally we analyzed the levels of each TF at MLL-AF4 peaks genome-wide, demonstrating the frequent colocalization of these proteins (Figure 3B, left). Strikingly, we also observed that as the width of the MLL-AF4 peak increases the width of the enriched TF signal correspondingly increases (Figure 3B, left), suggesting a direct relationship between the presence of MLL-AF4 and these TFs. To ask whether the reciprocal relationship was also true, we performed the same analysis at RUNX1 ChIP-seq peaks. Again, we observed increased ELF1, MAZ and MLL-AF4 signal, though there was no bias for MLL-AF4 spreading targets at stronger RUNX1 peaks (Figure 3B, right). To quantify the relationship specifically at the TSS of all genes, we compared MLL-N signal with RUNX1, ELF1 and MAZ signal and found correlations of 0.72, 0.67 and 0.69, respectively (Figure 3A).

**Figure 3.**
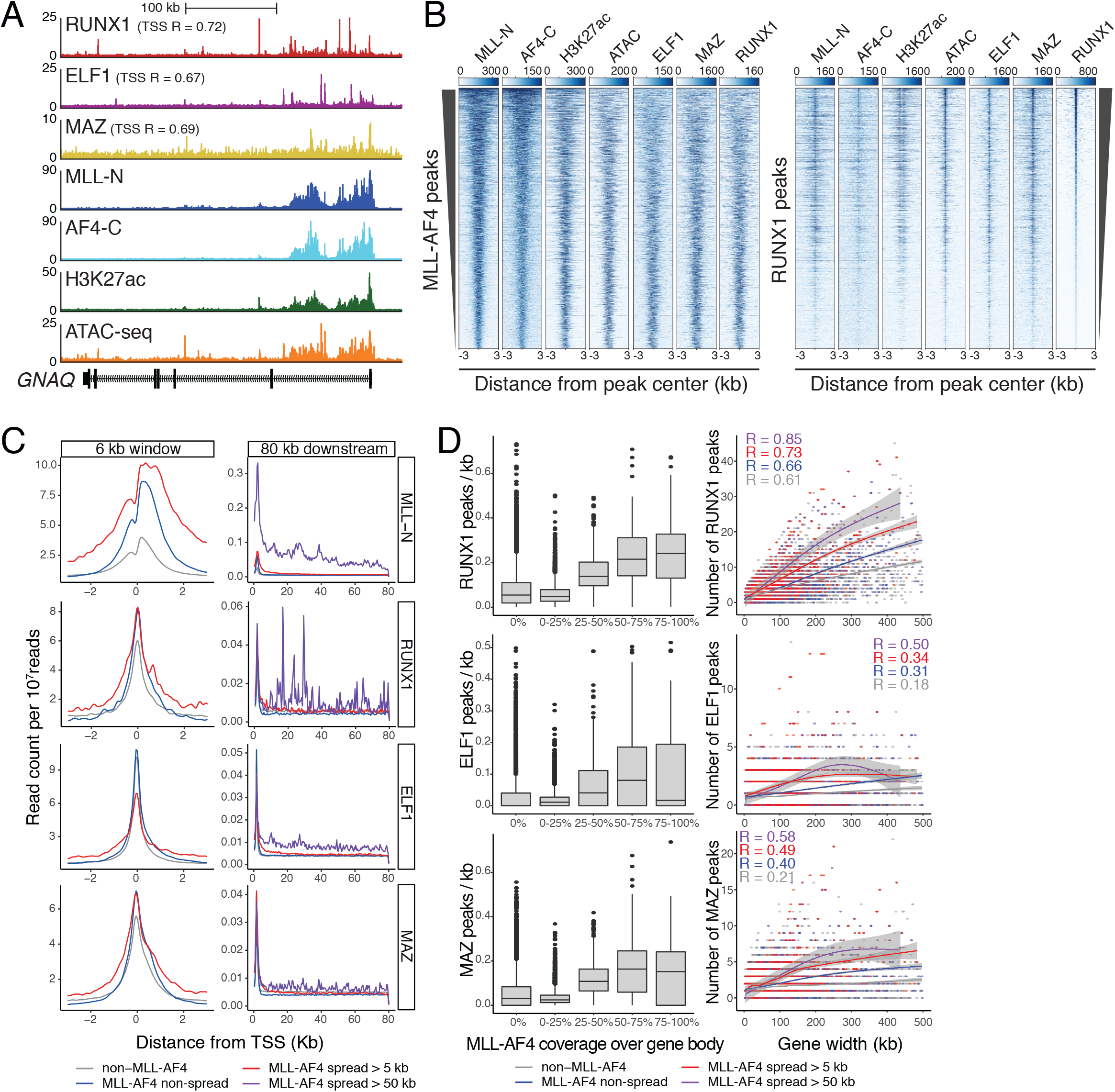
Core GRN factors are correlated with MLL-AF4 DNA binding profiles. (**A**) ChIP-seq tracks for MLL-N, AF4-C, RUNX1, ELF1, MAZ, H3K27ac and ATAC-seq over *GNAQ*. ChIP-seq data normalized to 1×10^7^ reads. Spearman’s rank correlations comparing RUNX1, ELF1 and MAZ signal with MLL-N signal over all TSS sites are shown. (**B**) Heatmaps of ChIP-seq datasets as well ATAC-seq in SEM cells. Left – Heatmaps generated over MLL-AF4 peaks with rows ordered on level of MLL-N and AF4-C reads. Right – Heatmaps generated over RUNX1 peaks with rows ordered on RUNX1 read count. (**C**) Meta-gene plots of ChIP-seq signal of expressed TSSs, from 3 kb up and downstream (left), or 80 kb downstream (right). Profiles shown include TSS not bound by MLL-AF4 (grey), MLL-AF4 bound TSS non-spreading (blue), MLL-AF4 bound TSS with spreading greater than 5 kb (red), and MLL-AF4 bound TSS spreading greater than 50 kb (purple). (**D**) Left – Boxplots showing number of TF ChIP-seq peaks/kb over gene bodies, grouped by percentage MLL-AF4 coverage of the gene body. Right – Scatter plot of number of TF ChIP-seq peaks in a gene body against gene body width (kb). Local regression lines (LOESS) describe the relationship between number of peaks and gene width, with grey ribbon representing 95% confidence intervals. Local regression lines calculated for genes with no MLL-AF4, MLL-AF4 non-spreading, MLL-AF4 spreading greater than 5 kb, and MLL-AF4 spreading greater than 50 kb.

To further investigate these correlations, we classified genes based on the extent of MLL-AF4 binding. Expressed genes were grouped, based on whether the TSS was not bound by MLL-AF4, only the TSS was bound, or where MLL-AF4 binding at the TSS spreads into the gene body by at least 5 kb (spreading targets). Based on these groupings, we created ChIP-seq meta-gene plots for each TF (Figure 3C, left). Both RUNX1 and MAZ showed an enrichment in signal directly over the TSS of MLL-AF4 bound targets but spreading targets did not show any further increase in signal. In contrast, ELF1 interestingly did not show a difference between non-MLL-AF4 and MLL-AF4 bound promoters, and even showed reduced signal at the TSS for spreading targets. However, TF levels away from the peak at the TSS were increased for all three factors at spreading targets, implying that while signal at the promoter is not strongly impacted in spreading targets, there is an increase in TF binding spreading into the gene body. To test this inference, we looked up to 80 kb into the gene body and added an additional subset of spreading targets with MLL-AF4-bound regions greater than 50 kb. Strikingly, at these genes we saw enrichment of all three TFs across the entire 80 kb window (Figure 3C, right).

This “spreading” of TF peaks may occur due to an increase in individual peak density, or an increase in peak width, as observed with MLL-AF4 and H3K27ac (Figure 3A). To assess these possibilities, we looked at peak density across active gene bodies. We categorized gene bodies by the percentage coverage of MLL-AF4, ranging from 0% (non-MLL-AF4 targets) to 100% (Figure 3D, left). We saw a clear increase in the number of TF peaks per kb of gene body correlated with increased MLL-AF4 coverage, suggesting that the higher frequency of TF binding events explains the enrichment at MLL-AF4 regions. To directly assess the effect of MLL-AF4 spreading, we plotted the total width of each gene body against the absolute number of intragenic peaks and stratified this by the MLL-AF4 binding profiles classified as in Figure 3C. As expected, we observed a correlation between the length of the gene body and the number of peaks for all three factors. What is more noteworthy is the slope of the relationship, which shows an increase in TF binding frequency at MLL-AF4 targets, and even greater at spreading targets, when comparing genes of the same length (Figure 3D, right). We also see a stronger correlation between gene length and TF peak count at spreading targets, highlighting a tighter relationship between these parameters. Of the three TFs, RUNX1 shows the clearest effect, and shows higher correlation, implying that it has a stronger relationship with MLL-AF4 binding. We tested whether this was a consequence of an increased frequency of RUNX1 binding motifs at MLL-AF4 spreading genes, but did not find a relationship between canonical RUNX1 motifs and MLL-AF4 spreading types. This implies that the TF enrichment may either be independent of the presence of DNA binding motifs, or that presence of MLL-AF4 allows for more stable binding of TF proteins to the motifs present (Supplementary Figure S3A). These analyses show that while MLL-AF4 and TFs do not necessarily have the same binding profiles, there is a clear increase in TF signal in regions marked with MLL-AF4. This appears to be through an increase in TF peak density across regions marked with the MLL-FP. As central TFs within the GRN, they are upregulated by MLL-AF4 and result in genome wide transcriptional changes beyond the scope of the fusion protein. Our analyses here suggest that MLL-AF4 also enhances the association of these factors with MLL-AF4 target genes, highlighting a positive feedback mechanism by which these key TFs further drive direct MLL-AF4 behaviors

### RUNX1 is a highly central and essential node of the MLL-AF4 network

Our analysis so far has identified several modules of GRN nodes displaying different relationships with patient leukemias, and a correlation between central GRN TFs and MLL-AF4 binding. To determine the functional importance of the key genes within the GRN we integrated data from published CRISPR essentiality screens. We initially extracted essentiality statistics from the Project Score database of the Cancer Dependency Map (DepMap), which has tested a large number of cancer cell line models (68). This database shows that MYC is pan-essential, with almost all cell lines displaying essentiality, and that MAZ is essential for none of the cell lines (Figure 4A). Interestingly, RUNX1 is only essential in two cell lines from the DepMap database. As we have previously identified RUNX1 as a key MLL-FP target in SEM cells (46) this was unexpected, though it is possible this screen lacks the statistical sensitivity for our purposes. However, one of the two cell lines, OPM-2, is a model for leukemia, specifically myeloma (88). We additionally used a published screen more specific to leukemia, involving two MLLr leukemia cell lines, MOLM-13 (MLL-AF9 AML) and MV4-11 (MLL-AF4 pediatric AML) (58). Using these, in combination with essentiality data from non-leukemia cancer cell lines HT-29 (colon adenocarcinoma) and HT-1080 (fibrosarcoma), we grouped the results of this screen into non-essential genes, non-leukemia essential, and essential specifically in either one or both leukemia cell lines (Figure 4B). For example, MYC is essential not only in leukemia cell lines, but also the non-leukemia cancer models, which implies it is generally essential for cancer cell function, and not specifically for leukemia in concordance with the DepMap database. Conversely, MAZ and ELF1, which are highly central in our MLL-AF4 GRN, were not found to be essential in any of these cell lines, suggesting that their roles in the network may be redundant with other proteins, or that their targets are not important for cell survival or proliferation, at least under the conditions of these CRISPR screens. It is important to note however that this is an *in vitro* screen, and that these non-essential TFs may well be essential for leukemia progression *in vivo*. In contrast to MAZ and ELF1, RUNX1 is a highly central factor that is specifically essential to both MOLM-13 and MV4-11 cell lines (Figure 4C, Supplementary Figure S4A). Other key leukemia-specific essential nodes here include MYB, MED13L, HOXA10, and the binding partner of RUNX1, CBFB. Considering that MAZ is inactive in normal FBM HSPCs, but appears to be central in MLL-AF4 leukemias, and that its binding correlates with MLL-AF4, we were surprised that it was not identified as functionally important for leukemia by either screen. We also found that knockdown of MAZ with siRNA showed minimal effects on the expression of key genes (Supplementary Figure S4B). This parallels the essentiality data and provides further evidence that although MAZ is a highly central node in the network, it is not a key TF required for leukemia cell survival, at least *in vitro*. Although MAZ may have little consequence on leukemia function, it could be a useful biomarker of MLL-FP activation.

**Figure 4.**
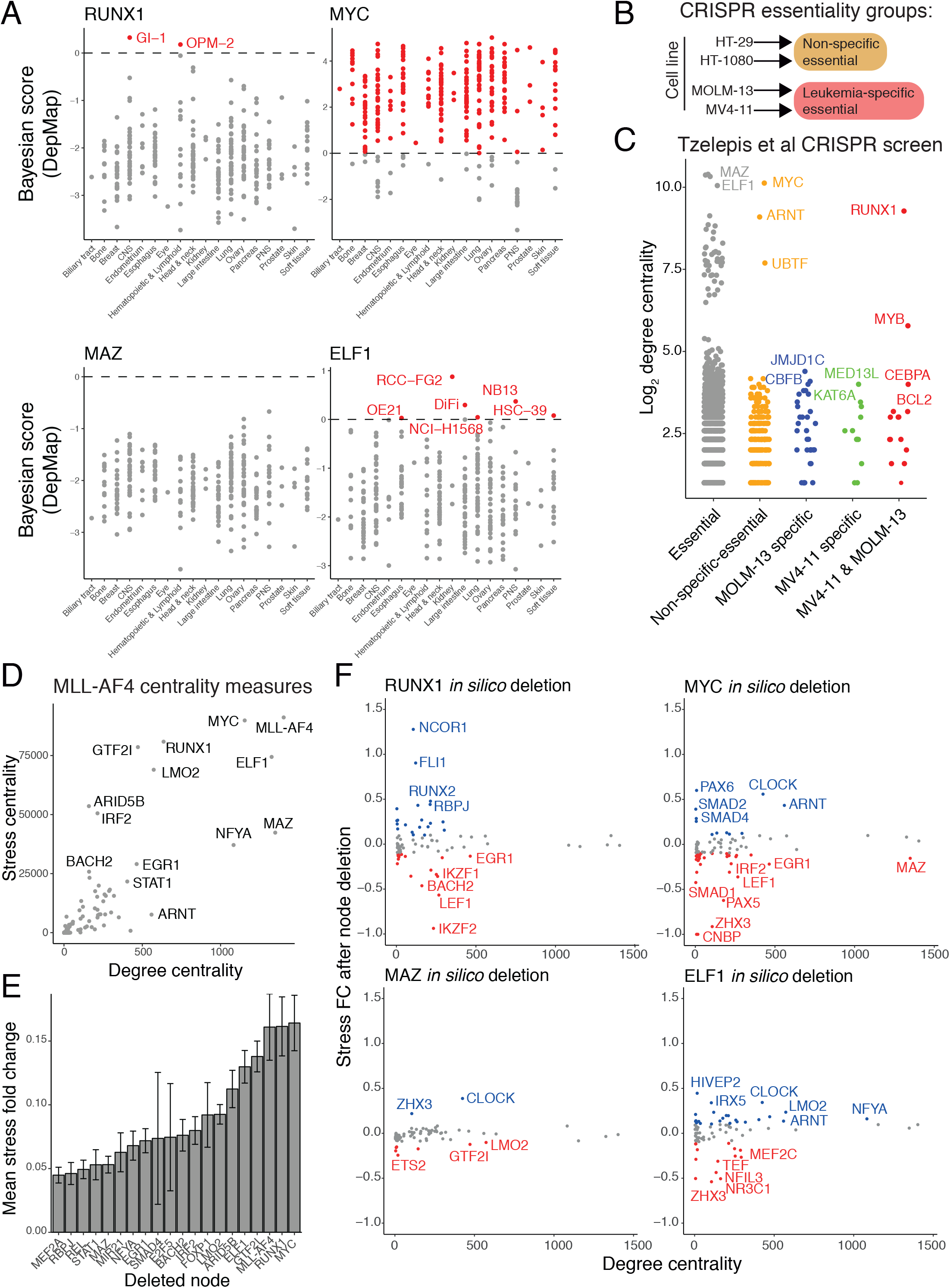
RUNX1 is a highly central and essential node in the MLL-AF4 network. (**A**) essentiality screen for RUNX1, MYC, MAZ and ELF1 from the Project Score database of the Cancer Dependency Map (68). Each datapoint represents a different cell line, with essentiality highlighted in red. Cell lines were categorized into the type of cancer that they model. Statistical analysis from the database is represented as Bayesian scores for which values above 0 indicate essentiality. (**B**) Schematic illustrating how the published CRISPR screen data (58) is used. Briefly, genes identified as essential in HT-29 (colon adenocarcinoma) or HT-1080 (Fibrosarcoma) were classified as non-specific essential. Genes not essential in HT-29 or HT-1080 but essential in MOLM-13 (MLL-AF9) or MV4-11 (MLL-AF4) were annotated as essential specifically for either MOLM-13 or MV4-11, or both cell lines. (**C**) Association of Log_2_ degree centrality with CRISPR essentiality groups as outlined in (**B**). Leukemia-specific essential genes were further categorized as specific to MOLM-13, MV4-11 or both. (**D**) Degree centrality plotted against stress centrality of SEM MLL-AF4 GRN nodes. (**E**) Mean absolute stress fold changes in response to *in silico* deletion of GRN nodes and subsequent recalculation of stress centrality. Stress fold change is calculated on a per gene basis in the GRN by contrasting stress scores before and after *in silico* deletion. Top 20 nodes by greatest stress fold change are shown. (**F**) Stress fold change responses for individual transcription factors plotted against degree centrality for *in silico* deletion of RUNX1, MAZ, ELF1 and MYC. Blue datapoints indicate positive stress fold change responses, while red indicate a negative response.

Stress centrality is a measure of the number of shortest paths between any other two points of a network that pass through a particular node (25). As there is a temporal delay between transcriptional activation of a gene and production of a TF protein, path length may reflect response times to perturbations (89). With this in mind, stress centrality could be an important metric for judging the importance of a node with respect to communication through the network. Within the MLL-AF4 GRN, stress and degree centrality are not perfectly correlated, with some highly connected but relatively low stress genes, such as MAZ (Figure 4D). Others are poorly connected with relatively high stress centrality, such as ARID5B and IRF2. ARID5B has been associated with IFN-γ production, and IRF2 is an interferon-stimulated gene (90, 91), which may imply that this high stress centrality reflects a specific role for these factors in the interferon response. RUNX1 has high stress centrality, though is not as well connected as nodes such as MYC or MAZ (Figure 4D). In order to predict the importance of a node for maintaining connectivity within the network, we performed *in silico* deletion of GRN nodes followed by recalculation of stress centrality. Loss of MLL-AF4, RUNX1 or MYC caused the greatest redistribution of stress centrality scores across nodes (Figure 4E). The implication may be that the shortest paths passing through these nodes are then rerouted through alternative genes. In particular, *in silico* deletion of RUNX1 led to increased NCOR1 and FLI1 stress, indicating these may act as alternative intermediate nodes to RUNX1 (Figure 4F, Supplementary Figure S4C). These data strongly point towards RUNX1 as a key node in the MLL-AF4 GRN.

### RUNX1 and MLL-AF4 regulate targets in feed-forward loops and cascade motifs

We have previously established that MLL-AF4 induces overexpression of RUNX1 (46). As RUNX1 appears to be an important node, both in terms of its centrality within the MLL-AF4 GRN and its essentiality in leukemia cell lines, we next wanted to assess the impact of RUNX1 disruption on gene expression in SEM cells. We performed siRNA knockdown of RUNX1 for 96 hours followed by nascent RNA-seq and identified 13914 expressed genes, 2662 of which were upregulated and 2550 downregulated (Supplementary Data S3). The majority of these DEGs were bound by RUNX1 (89% and 88% respectively) (Figure 5A, Supplementary Figure S5A-B). We compared the DEGs following RUNX1 knockdown with MLL-AF4 knockdown to identify genes that were dysregulated by the loss of either protein. There were 2279 genes affected by both treatments, with 2933 and 2519 genes specific to RUNX1 and MLL-AF4 knockdown, respectively (Figure 5B). Common DEGs were significantly enriched for GO terms related to hematopoiesis, regulation of cell death, and positive regulation of B cell proliferation, consistent with a leukemic expression program (Figure 5C). Additionally, we saw enrichment for pathways related to DNA methylation, general epigenetic regulation of gene expression, and cellular senescence (Supplementary Figure S5C). To further integrate RUNX1 regulatory behavior with MLL-AF4 we created a RUNX1-centric GRN using the same methodology as outlined in Figure 1D, by integrating RUNX1 knockdown RNA-seq and ChIP-seq (Supplementary Data S1). The most central nodes of this network include established targets of RUNX1 regulation such as GFI1 (92), and MYB (93) (Figure 5D). As we propose that MLL-AF4 regulates its network partly through regulation of intermediate TFs, including RUNX1, we would expect that the RUNX1 GRN should have significant overlap with the MLL-AF4 GRN. To appropriately gauge the extent of overlap we also compared them with the BRD4 and DOT1L GRNs (Supplementary Figure S1B-C). As previously discussed, the DOT1L GRN overlaps with greater specificity to MLL-AF4, while BRD4 has a much more general regulatory scope. The RUNX1 GRN itself overlaps with a greater proportion of the MLL-AF4 GRN than DOT1L (1930 genes for RUNX1, 1363 for DOT1L), and shows a greater significance of overlap than both DOT1L and BRD4 (Supplementary Figure S5D-E). RUNX1 appears to have an overlapping role with MLL-AF4 that may drive leukemogenesis and appears to more specifically target the MLL-AF4 transcriptional network than IBET or DOT1Li, suggesting that RUNX1 may be responsible for a significant proportion of the direct and indirect transcriptional effects of MLL-AF4.

**Figure 5.**
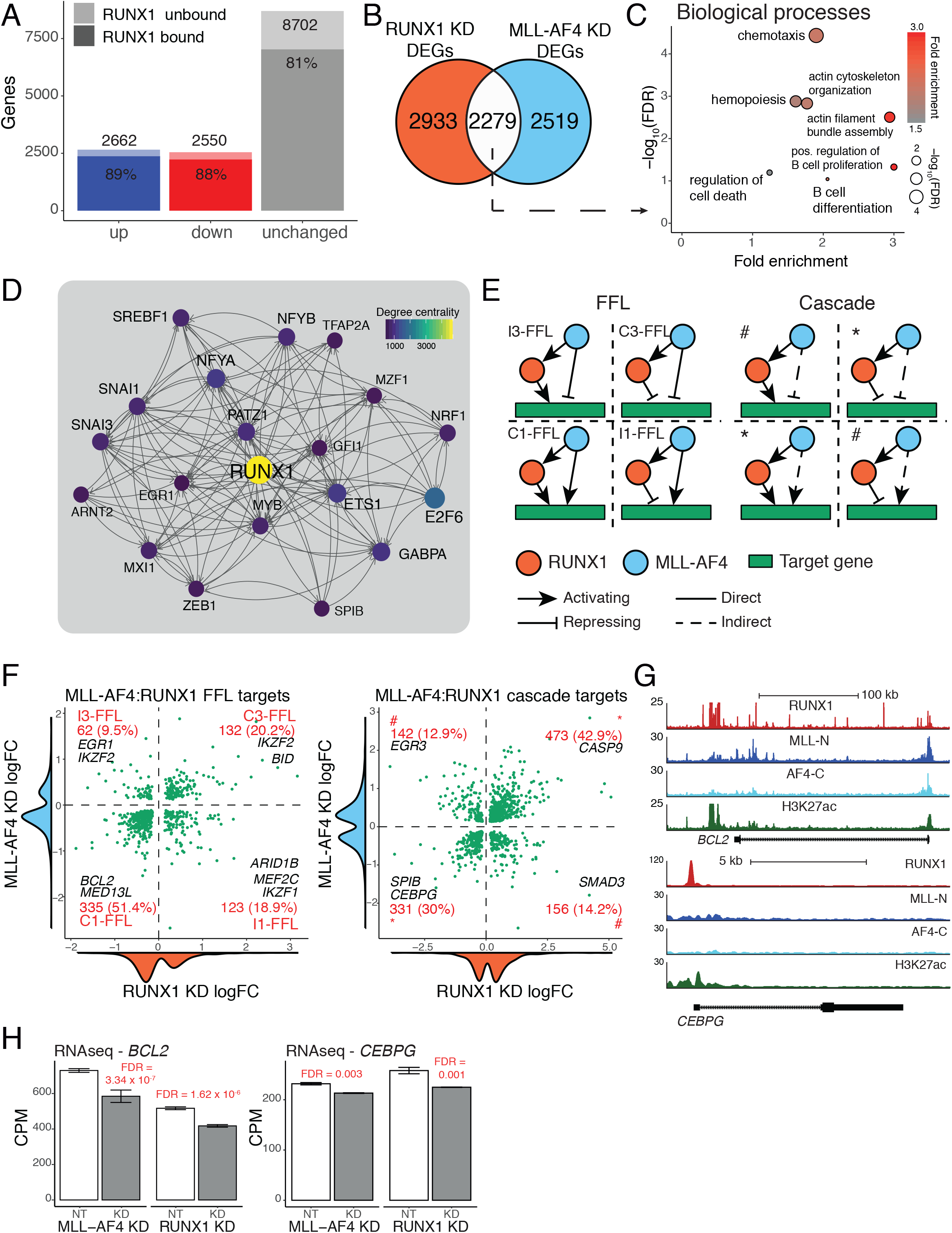
MLL-AF4 cooperates with RUNX1 in FFL and cascade circuits to regulate downstream targets. (**A**) DEGs from nascent RNA-seq after 96 hours’ RUNX1 siRNA knockdown. Differential expression is defined as FDR < 0.05. (n=3). Shaded area represents genes bound by RUNX1. (**B**) Venn diagram showing overlap between MLL-AF4 knockdown DEGs (Figure 1C) and RUNX1 knockdown DEGs (**A**). (**C**) GO biological process enrichment for overlap between MLL-AF4 and RUNX1 knockdown DEGs (**B**). Size of points is proportional to the significance of the enrichment, while point color represents fold enrichment over expected number of genes. (**D**) Top 20 genes of the RUNX1 GRN by degree centrality. Lines (edges) indicate predicted interaction from protein (source) to gene locus (target), with arrowheads pointing towards downstream nodes. (**E**) A schematic showing potential FFL (left) and cascade (right) motifs. FFLs are categorized into subtypes including coherent type 1 (C1-FFL), coherent type 3 (C3-FFL), incoherent type 1 (I1-FFL) and incoherent type 3 (I3-FFL). Solid lines represent direct interactions, dashed lines represent indirect. Cascade motifs are grouped into same sign of effect (*) and opposing sign of effect (#). (**F**) Scatter plot of FFL (left) and cascade (right) motif target genes displaying logFC response to RUNX1 knockdown and MLL-AF4 knockdown on the x and y axis, respectively. Along each axis is a density plot illustrating the distribution of logFC values. −logFC values are assumed to imply the perturbed factor has a positive regulatory effect on the target, while +logFC values are assumed to imply a negative regulatory effect. Quadrants of scatter plots align with FFL and cascade types shown in (**E**). (**G**) ChIP-seq tracks generated in hg19 for MLL-N, AF4-C, RUNX1 and H3K27ac. ChIP-seq data normalized to 1×10^7^ reads. Top locus is for *BCL2*, an example of an FFL target, bound by both MLL-AF4 and RUNX1. Bottom locus is *CEBPG*, an example of a cascade target, bound by RUNX1 but not MLL-AF4. (**H**) Expression of *BCL2* (left) and *CEBPG* (right) from nascent RNA-seq data following knockdowns. Expression normalized as CPM.

To assess how RUNX1 behaves in the context of the MLL-AF4 network, we took advantage of the functional RNA-seq and ChIP-seq data used for both models and integrated the two GRNs together. This allowed us to better understand the details of how these two regulatory proteins interact at individual gene targets and gave greater confidence to observed interactions. We then abstracted the resulting GRN into a series of three-node motifs, in order to understand the interaction between MLL-AF4 and RUNX1 activity. To analyze the combined MLL-AF4/RUNX1 GRN, we extracted gene targets that are regulated in specific three-node motifs involving both RUNX1 and MLL-AF4 (Figure 5E, Supplementary Data S4). We focused on FFLs, as these are prevalent in GRNs (9), and cascades, as they may mediate indirect effects of MLL-AF4. As we have previously established RUNX1 as a target of MLL-AF4 (46), a FFL here can be described as a motif where MLL-AF4 regulates RUNX1, and both MLL-AF4 and RUNX1 co-regulate a gene target (Figure 5E). A cascade can be described as MLL-AF4 regulating RUNX1, and RUNX1 regulating a gene target which is not bound by MLL-AF4 (Figure 5E). FFLs can be further grouped into coherent and incoherent types, based on the regulatory effect on the target gene from both RUNX1 and MLL-AF4, where coherent refers to same sign of effect and incoherent refers to opposing signs. This configuration can be broken into eight different types of FFL based on the combination of positive or negative interaction signs, as described by the Alon lab (13). As we know RUNX1 is regulated positively by MLL-AF4, this provides us with only four types to account for, type 1 and 3 coherent (C1-FFL, C3-FFL) and type 1 and 3 incoherent FFLs (I1-FFL, I3-FFL) (Figure 5E). We used the knockdown RNA-seq data to infer positive or negative effects on a gene target, with the assumption that changes are a consequence of direct interaction from MLL-AF4 and RUNX1.

We identified 652 potential FFL targets co-regulated by MLL-AF4 and RUNX1, 71.6% of which were coherent, suggesting that MLL-AF4 and RUNX1 predominantly act cooperatively (Figure 5F, left). C1-FFLs, where both TFs positively regulate a target, have previously been shown to be the most common type of FFL in other systems (11, 12), which is recapitulated in our data where 51.4% of targets are positively regulated by RUNX1 and MLL-AF4. We also identified 1102 potential cascade targets, which are bound by RUNX1 but not MLL-AF4. The majority of predicted cascade targets show an agreement in transcriptional response to MLL-AF4 and RUNX1 knockdown, showing common activation (331, 30%) or repression (473, 42.9%) (Figure 5F, right). This is in line with our hypothesis that indirect transcriptional changes from MLL-AF4 knockdowns are mediated by the activity of downstream TFs such as RUNX1. Note that “incoherent” target gene responses for cascade interactions, as illustrated in Figure 5E (denoted with #), are likely due to the complication of additional TFs regulated by MLL-AF4 that function at these sites. These are genes that appear to be activated by RUNX1 and repressed by MLL-AF4, or repressed by RUNX1 and activated by MLL-AF4 (142, 12.9% and 156, 14.2%, respectively).

An interesting observation from these analyses is that FFLs show a bias towards synergistic upregulation of target genes, while target gene regulation by cascade motifs is more balanced between activation and repression. This is congruent with the hypothesis that MLL-AF4 functions through promoting transcription and suggests that MLL-AF4 can bias RUNX1 activity towards promoting gene expression. For example, the known MLL-AF4 target *BCL2* (94), which is an anti-apoptotic factor, is activated in a C1-FFL (Figure 5G-H). Conversely, the pro-apoptotic factor *BID* is repressed by both TFs in a C3-FFL (Supplementary Figure S5F). Overall, it is striking that the majority of targets bound by RUNX1, and indirectly regulated by MLL-AF4, display the same regulatory logic after knockdown of RUNX1 or MLL-AF4 (72.9% of targets). One example of this is an activating cascade targeting the *CEBPG* locus, which is bound by RUNX1 but not MLL-AF4, and shows decreased transcription upon knockdown of either factor (Figure 5G-H). This suggests that RUNX1 binding and activity is a strong determining factor for the regulation of cascade gene targets and that RUNX1 is capable of mediating both activation and repression within the same regulatory network. This is consistent with published observations of RUNX1 activity in hematopoietic differentiation (95, 96).

### A cascade circuit regulates apoptosis through repression of *CASP9*

Having demonstrated that RUNX1 is a key node in the MLL-AF4 network, and identified FFL and cascade regulatory patterns controlled through the MLL-AF4:RUNX1 interaction, we wanted to test whether these structures had physiological relevance. To address this, we took a phenotypic screening approach. We wanted to focus on the regulation of apoptosis, as this process is affected by both MLL-AF4 and RUNX1 siRNA knockdowns (Figure 5C), and particularly since MLL-FP leukemias are highly sensitive to induction of the BCL-2 apoptosis pathway (50, 55, 56). This is due in part to the fact that MLL-AF4 directly regulates key genes in the BCL-2 apoptosis pathway (50, 94), but little is known about how a wider GRN could control this pathway.

We found that RUNX1 is predicted to regulate 30 components of the apoptosis KEGG pathway (hsa04210), compared to 20 regulated by MLL-AF4 (Figure 6A). These predicted interactions do not necessarily prescribe pro- or anti-apoptotic results. To describe in detail how TFs of the GRN interact with the apoptosis pathway we created an interaction matrix (Figure 6B, Supplementary Figure S6A), which functions as a set of hypotheses for regulation of apoptosis. It is worth noting that each TF of the GRN regulates a different but overlapping set of apoptosis-related genes, which highlights how no single TF can provide a complete picture describing regulation of the pathway. *BCL2* is predicted to be regulated by many different factors, including MLL-AF4, RUNX1, ELF1 and MYC (Figure 6B). This TF cooperation may allow for strong and robust regulation of *BCL2* that is resistant to upstream perturbations and may help to explain why *BCL2* is highly expressed in MLL-AF4 leukemias (50, 97). A number of genes in the apoptosis pathway are regulated by MLL-AF4 and/or RUNX1. For example, FFLs regulate *CASP8, BID* and *BCL2*, whereas *APAF1* and *CASP9* are regulated in cascades, and *CASP3* and *CASP7* are targeted directly by MLL-AF4 independently of RUNX1 (Figure 6B).

**Figure 6.**
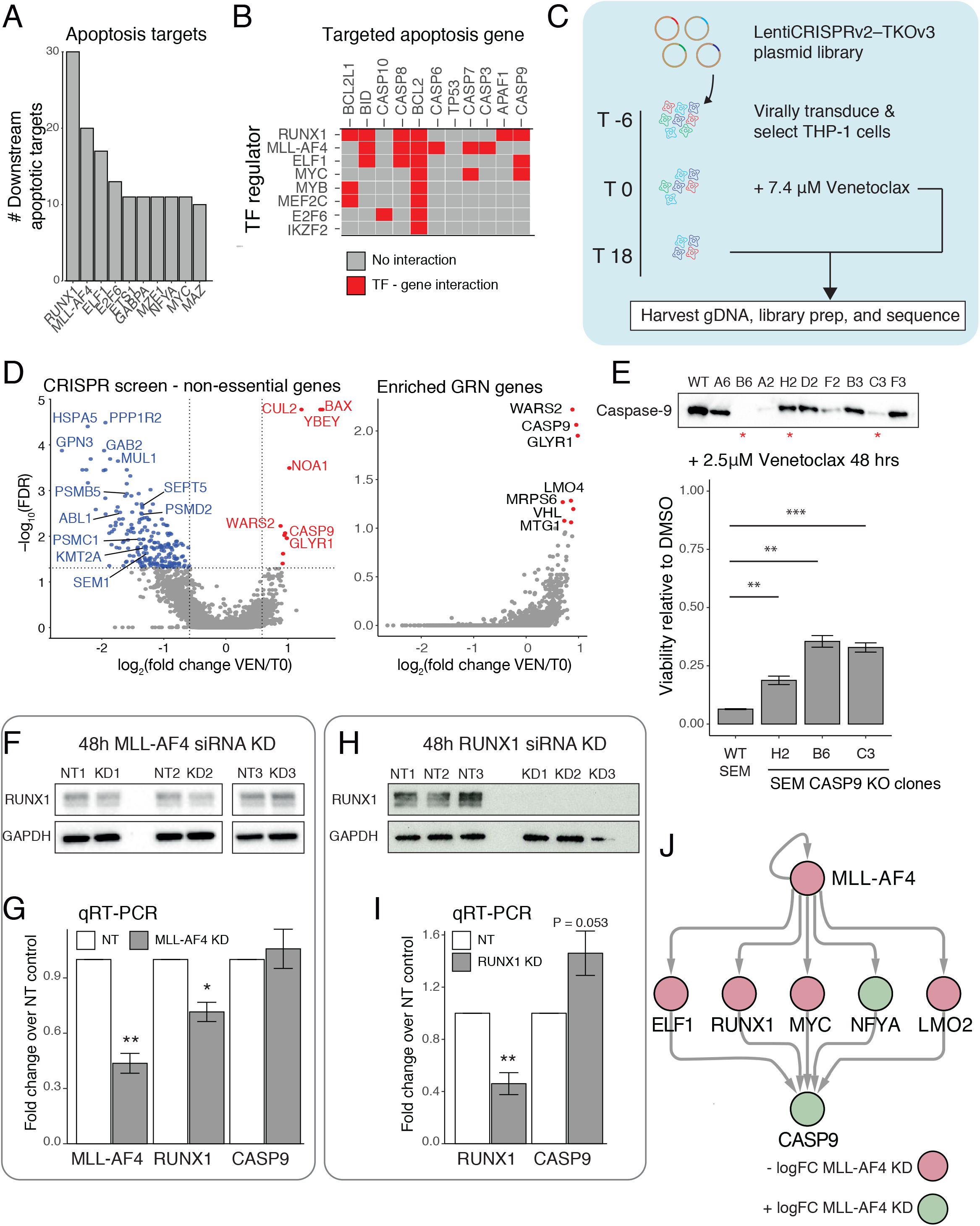
*CASP9* is a key apoptotic gene repressed by MLL-AF4 in a cascade motif. (**A**) Number of apoptosis KEGG pathway (hsa04210) components regulated by TFs within the MLL-AF4 GRN. Top 10 apoptosis regulators are shown. (**B**) Interaction matrix showing a subset of TF regulators of apoptosis pathway (rows) and components of the apoptosis pathway (columns). Red squares indicate a predicted regulatory interaction between the TF and apoptosis component. Full interaction matrix shown in Supplementary Figure S6A. (**C**) Experimental outline of CRISPR screen performed in combination with 7.4 μM venetoclax treatment of THP-1 cells. Cells harvested for sgRNA counting and analysis at T0 (after transduction and selection) and T18 (18 days treatment with venetoclax). (**D**) Analysis of CRISPR screen using MAGeCK, plotting −log_10_ FDR against log_2_ gRNA fold change. Left – MAGeCK results, excluding pan-essential genes. Right – MAGeCK results for enriched sgRNA, filtered for MLL-AF4 GRN nodes. (**E**) Top – Western blot for Caspase-9 from *CASP9* knockout clones in SEM cells. Clones used for functional analysis are indicated by *. Bottom – CellTiter-Glo viability assay of CASP9 knockout clones after 48 hours treatment with 2.5 μM venetoclax. CTG luminescence displayed as a ratio relative to DMSO control. (n=3, ** < 0.01, *** < 0.001). Error bars represent standard error of the mean. (**F**) Western blot in SEM cells showing RUNX1 protein levels after 48 hours’ MLL-AF4 knockdown. GAPDH used as a loading control. (**G**) qRT-PCR results showing expression of *MLL-AF4*, *RUNX1* and *CASP9* following 48 hours’ MLL-AF4 siRNA knockdown (n=3, * < 0.05, ** < 0.01). Error bars represent standard error of the mean. (**H**) Western blot in SEM cells showing RUNX1 protein levels after 48 hours’ RUNX1 knockdown. GAPDH used as a loading control. (**I**) qRT-PCR results showing expression of *RUNX1* and *CASP9* following 48 hours’ RUNX1 siRNA knockdown (n=3, ** < 0.01). Error bars represent standard error of the mean. (**J**) Illustration of selected cascade motifs within the MLL-AF4 GRN that explain regulatory connectivity between MLL-AF4 and CASP9. Node color represents logFC response to MLL-AF4 knockdown.

To screen these targets for their importance in regulating cell death under potential therapeutic conditions, we made use of venetoclax, a BCL-2 inhibitor drug that is a promising treatment for MLLr ALLs (50, 55, 56). We performed a CRISPR screen with the TKOv3 genome-wide sgRNA library (69, 70) in the presence of venetoclax, to drive apoptosis and detect gene perturbations that increased, or decreased, cell survival from drug induced apoptosis (Figure 6C). We opted to use THP-1 cells, as they are an MLL-AF9 driven model of AML that is only moderately sensitive to venetoclax and can be used more easily to screen for both induction of resistance as well as sensitivity. We reasoned that since core aspects of the MLL-AF4 GRN are highly conserved more generally in AML and ALL (see Figure 2), core regulatory circuits of the GRN should be conserved in AML cells. TKOv3 CRISPR KO cells were treated with 7.4 μM venetoclax for 18 days, and NGS was used to compare sgRNA frequency in the pool before (T0) and after drug treatment (VEN). We used the Annexin-V/PI apoptosis assay to determine that 7.4 μM is the half-maximal cytotoxic concentration of venetoclax in THP-1 cells (Supplementary Figure S6B). Our sgRNA pool showed reliable representation of all sgRNAs (Supplementary Figure S6C). Plotting sgRNA counts between biological replicates showed good reproducibility of the screen (Supplementary Figure S6D). Analysis of enriched and depleted sgRNA (Supplementary Data S5), excluding known pan-essential genes (Supplementary Figure S6E), resulted in many hits (Figure 6D, left, Supplementary Figure S6I). Depletion of a sgRNA indicates that loss of the targeted gene promotes cell death in combination with venetoclax, whereas enrichment of a sgRNA suggests that gene knockout attenuates the effect of venetoclax. *KMT2A* targeting sgRNA, which would result in inactivation of the essential MLL-AF9 fusion protein, is shown to be depleted after venetoclax treatment. We also observed depletion of sgRNA for proteasome subunits (PSMB5, PSMC1, PSMD2), suggesting an essential role for this complex in cell survival. Enrichment of sgRNAs targeting the pro-apoptotic BAX are consistent with the inhibition of BAX by BCL-2, which is relieved by venetoclax. We validated select results from the screen by assaying for cell viability following transduction with individual sgRNA (Supplementary Figure S6F). We used sgRNA sequences for validation from the Brunello library (98) to ensure the results were not due to off-target effects of the TKOv3 counterparts. Focusing specifically on enriched sgRNA that are present in the GRN (Figure 6B), we identified *CASP9* as one of the most significant results (Figure 6D, right). This implies that deletion of *CASP9* promotes survival in the face of venetoclax-mediated apoptosis, consistent with the activation of Caspase-9, and subsequent signaling cascades, following the release of cytochrome c from mitochondria, the process inhibited by BCL-2 (99–101).

The venetoclax/CRISPR screen in THP-1 cells identified Caspase-9 as a potentially important component of the BCL-2 apoptosis pathway in these cells. Following from our rationale that key components of the GRN are conserved between AML and ALL, we wanted to determine if Caspase-9 is important in MLL-AF4 ALL cells and if *CASP9* is regulated by the MLL-AF4 GRN. To confirm the importance of Caspase-9 in MLL-AF4 ALL cells we used CRISPR sgRNA from the Brunello library to stably delete *CASP9* in SEM cells (Figure 6E). We selected two knockout lines with no detectable Caspase-9 protein (B6, C3), and a line displaying partial loss of protein (H2). We treated cells with either DMSO or 2.5 μM venetoclax for 48 hours and tested cell viability using CellTiter Glo (CTG) assays (Figure 6E). We observed a significant increase in viability for both the partial knockout in H2 (P = 0.0061), and the complete knockout lines, B6 and C3 (P = 0.0013, P= 0.0008 respectively) (Figure 6E), confirming that *CASP9* is important for venetoclax-induced cell death. Based on our matrix of interactions (Figure 6B) we predicted that an MLL-AF4 and RUNX1 cascade regulates *CASP9,* as the gene is bound by RUNX1 and not MLL-AF4 (Supplementary Figure S6G. Both MLL-AF4 knockdown and RUNX1 knockdown leads to an increase in *CASP9* expression, which implies that it is repressed by these factors (Supplementary Figure S6H).

The MLL-AF4 knockdown data used covers 96 hours’ siRNA treatment, at which point RUNX1 expression is impacted. It was therefore unclear whether the effect of MLL-AF4 knockdown on *CASP9* was a consequence of MLL-AF4 loss or an indirect consequence of *RUNX1* downregulation. To test these possibilities, we performed the *MLL-AF4* siRNA treatment for 48 hours, at which point RUNX1 was only minimally affected at the protein level (Figure 6F). We confirmed a reduction in *MLL-AF4* mRNA levels by qRT-PCR (Figure 6G), and, consistent with its role as a direct target of MLL-AF4, we observed some reduced *RUNX1* expression, although protein levels were not yet affected (Figure 6F). Importantly, we did not see an effect on *CASP9* expression at this time point (Figure 6G, P = 0.64). In contrast, after siRNA knockdown of *RUNX1* for 48 hours, RUNX1 protein was undetectable (Figure 6H), and there was a significant reduction in *RUNX1* mRNA levels (Figure 6I). In contrast to MLL-AF4 knockdown, we also observed an increase in *CASP9* expression (Figure 6I, P = 0.053). Together these data show that MLL-AF4 disruption on its own is not sufficient to significantly affect *CASP9* expression, but instead it acts via RUNX1 to repress the gene. There are likely other TFs that also play an intermediate role in this circuit alongside RUNX1, and our GRN network proposes several possible factors (Figure 6J). This leads to the proposed gene regulatory pathway in which MLL-AF4 acts with downstream TFs, such as RUNX1, in a cascade fashion to suppress *CASP9* expression, and therefore inhibit apoptosis. Together these results highlight the utility of using a GRN to predict and analyze the wide-ranging regulatory consequences of MLL-AF4 activity, and the complex ways in which both repression and activation of key gene targets are maintained in these leukemias (Figure 7).

**Figure 7.**
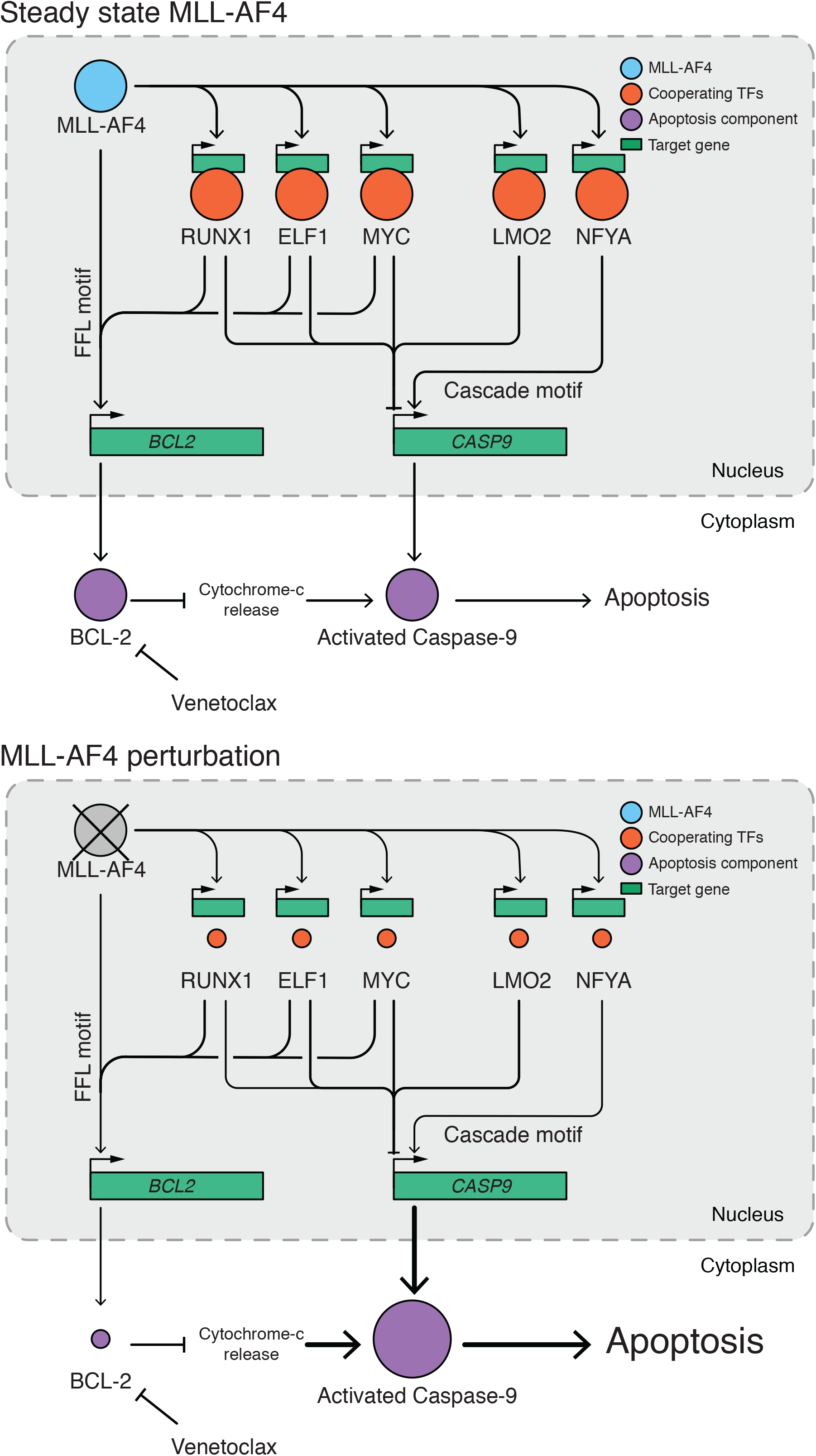
A model of MLL-AF4 driven FFL and cascade network motifs that interface with the apoptosis pathway. *BCL2* is regulated through a number of FFL motifs involving a range of TFs, which provides robust activation of the locus. *CASP9* instead is regulated through cascade motifs. Many of the TFs share a role in FFL activation of *BCL2* and cascade repression of *CASP9*. This dual function of TFs allows for more broad regulation of apoptosis than the direct behavior of the MLL-AF4 fusion protein alone allows. We use this model to propose that in the absence of MLL-AF4 these activated TFs decrease in expression, resulting in a marked loss of *BCL2* activating signal, and a loss of *CASP9* repression, shifting the balance towards apoptosis. As cancers are highly adaptable and prone to drug resistance, these cooperating TFs are avenues by which cancer cells may mediate resistance mechanisms that rebalance the signals feeding in to *BCL2* and *CASP9*, through mutagenesis or genome wide epigenetic changes.

## DISCUSSION

Knockdown of MLL-AF4 resulted in both up- and downregulation of transcription, and unexpectedly, the majority of these genes did not involve direct MLL-FP binding. This observation poses the question: if MLL-AF4 predominantly functions directly through promoting transcriptional activation, by what mechanisms can it regulate gene activation and repression in a wider network?

In this study we have created a GRN that specifically probes the relationship between MLL-AF4, the primary driver of leukemic transformation (47–49), and the downstream transcriptional network susceptible to perturbation of the fusion protein. Our systematic approach to explain indirect MLL-AF4 interactions led to the establishment of a range of predicted circuits, including the fact that a key downstream TF in the MLL-AF4 GRN includes RUNX1. Although the importance of RUNX1 in MLL-FP leukemias has been observed previously (46, 102), placing RUNX1 in the context of the MLL-AF4 GRN allows us to capture complex aspects of the MLL-AF4:RUNX1 regulatory axis, highlighted by our observation of the shared regulatory logic of RUNX1 and MLL-AF4 at the majority of gene targets.

In addition, we are also able to make predictions about how the GRN regulates cell behavior through MLL-AF4:TF controlled network motifs. For example, to find circuits relevant to leukemia we screened for targets that modulate response to venetoclax-induced apoptosis. With this approach we identified an MLL-AF4:RUNX1:*CASP9* repressive cascade, which may prevent cell death in tandem with FFL motifs activating *BCL2* (Figure 7). This suggests a potential mechanism for MLLr cells to acquire resistance to venetoclax; as enhanced activity of the MLL-AF4:RUNX1:*CASP9* cascade would result in stronger repression of *CASP9* and therefore reduced apoptosis. However, our GRN model proposes a number of intermediate factors other than RUNX1 that may regulate *CASP9*, and thus we can expand our detailed understanding of this regulatory axis by considering the combinatorial effect of multiple MLL-AF4:TF:*CASP9* cascade motifs. With this in mind, we can conclude that cascade motifs can promote cell survival, although more work is required to determine the extent of this effect in a stable leukemia state. However, the successful detection of a cascade motif using our GRN that explains the wider transcriptional control of MLL-AF4, and the systematic approach of motif screening is a strong proof of principle and highlights the importance of not only studying the direct effect of MLL-FP behaviors, but also assessing the overall transcriptomic context.

### MLL-AF4 driven network motifs are important for understanding MLL-AF4 fusion protein behaviour

Our current study identifies a number of cascade and FFL motifs and validates a key TF cascade (MLL-AF4:RUNX1:*CASP9*), which offers a proof of principle for MLL-AF4 engagement of a wider transcriptional network. We found that FFLs, which are generally enriched in biological networks (9), may also be important for synergistic enhancement of core MLL-AF4 activity. Central TFs in our model correlate with MLL-AF4 DNA binding, leading to a high occurrence of FFL motifs. We have not yet elucidated the mechanism behind this correlation, or if it is functionally relevant, though we have found this is not due to an increased frequency of canonical RUNX1 DNA binding motifs within these genes. It may be that MLL-AF4 binding creates a permissive environment that encourages TF binding, or that high TF binding density encourages MLL-AF4 binding or spreading; or the two processes together may form a positive feedback loop for enhanced binding of MLL-AF4 and TFs. We also highlight an interplay between different types of regulatory motifs, as the MLL-AF4:TF:*CASP9* repressive cascades identified here may cooperate with FFLs activating *BCL2* thus preventing cell death (Figure 7). This provides a potentially useful paradigm where simple patterns of cascade and FFL motifs can cooperate in regulating complex leukemia promoting pathways.

### Circuits that describe the GRN model may function as the basis for understanding leukemic behavior

One potential caveat of our approach is that it is highly dependent on cells grown in culture. It has been established that immortalized cell lines show considerable changes in the regulatory landscape when compared with primary tissues. Transcription in cell lines still reflects original tissue behaviors, but particular pathways are strongly impacted, such as the cell cycle (77). Indeed, integrating our GRN with patient data did reveal a cell line-specific cluster of nodes, that is enriched for cell cycle and stress response terms. Identification of this cluster allowed us to separate out these clinically irrelevant factors, and instead focus on key pathways that are conserved in the patient data. Ultimately the real test of any set of key targets for relevance to leukemia is through the analysis of patient samples derived from clinical trials, in which GRN identified pathways may be therapeutically targeted in an *in vivo* context. The main advantage of our network is that it provides a framework for analyzing datasets that could otherwise be overwhelmingly complex.

An interesting concept briefly discussed in a review from the Pimanda lab (3) is that AML is caused not by a particular mutation but instead by dysregulation of the transcriptional network. This would explain why we observe such a wide range of chromatin/transcription-associated driver mutations that may function through disruption of the same transcriptional circuits. Our data reinforces this concept as we see that the most connected TFs of the GRN are ubiquitously expressed across different ALL and AML patients. This may also help to explain why MLL-AF4 is sufficient to drive leukemogenesis in the absence of other cooperating mutations. The development of a cancer cell requires the disruption of multiple processes, usually necessitating multiple cooperative mutation events (103). However, the MLL-AF4 network on its own may be able to co-opt multiple pathways to achieve a similar result. One would then expect many of these circuit connections to be conserved in other leukemias, as they could represent common regulatory wiring that enables an oncogenic transcription program, but that are driven by a variety of other initiating events. The fact that ALL MLLr leukemias have been known to switch to an AML lineage (33, 34) further implies that core MLLr driven circuits attribute to multiple types of leukemia. MAZ is the most striking example of a central, leukemia specific factor, that is not expressed in normal FBM cells. A recent study has identified MAZ in having a role in the TF network that describes embryonic HSC ontogeny (87), which may indicate that circuits related to embryonic hematopoiesis are co-opted in leukemia.

Future GRN studies, that extensively characterize multiple types of leukemia, may allow us to identify common, core FFL and cascade motifs that are generated by multiple leukemic drivers and characterize myeloid and lymphoid leukemias. Comparative analyses using ATAC-seq to determine differential cis-regulatory elements, such as is performed in an AML sub-type study from the Bonifer lab (23), would be particularly insightful for determining core and type-specific leukemic circuits. The identification of core regulatory wiring common to all leukemias would improve our understanding of leukemia progression, and more importantly, this could lead to development of therapies designed to target these core pathways.

### FFL and cascade motifs can suggest mechanisms by which regulatory signals may rebalance to establish drug resistance

A key concept for drug resistance acquisition is that cancers can exhibit changes in epigenetic regulation which may rebalance regulatory wiring (104). Knowledge of GRN circuits may identify mechanisms by which epigenetic changes may alter apoptosis components such as *CASP9*. For example, adaptations that increase *RUNX1* expression could result in increased *BCL2* and reduced *CASP9* mRNA levels (Figure 7). This raises the possibility that combined treatment with RUNX1 inhibitors could be an effective therapeutic strategy. We also identify a number of other cooperative TFs that may regulate *CASP9* and *BCL2,* showing that the GRN can be used to highlight predictions in ways which compensation of a robust pathway may occur.

This capacity for adaptation, using existing pathways, allows us to use these regulatory motifs to suggest mechanisms by which drug resistance may occur. The concept of epigenetic rebalancing of regulatory circuits is reinforced by previous work showing that targeting epigenetic marks, such as H3K79me3 through DOT1Li, synergizes with venetoclax treatment (50, 105). One hope is that understanding the wider GRN of leukemias will allow for the design of combinatorial therapies that could help prevent the acquisition of drug resistance after treatment.

## ACCESSION NUMBERS

Previously published sequencing data are available from GEO accession numbers reported in Supplementary Table S2. RNA-seq and ChIP-seq data generated in this study are available under GSExxxxxx.

## SUPPLEMENTARY DATA

Supplementary Data are available at NAR online.

## ACKNOWLEDGEMENT

We thank the MRC WIMM Centre for Computational Biology (CCB), Radcliffe Department of Medicine, University of Oxford; and Jelena Telenius for the use of her pipelines.

---

**Supplementary Figure 1.**
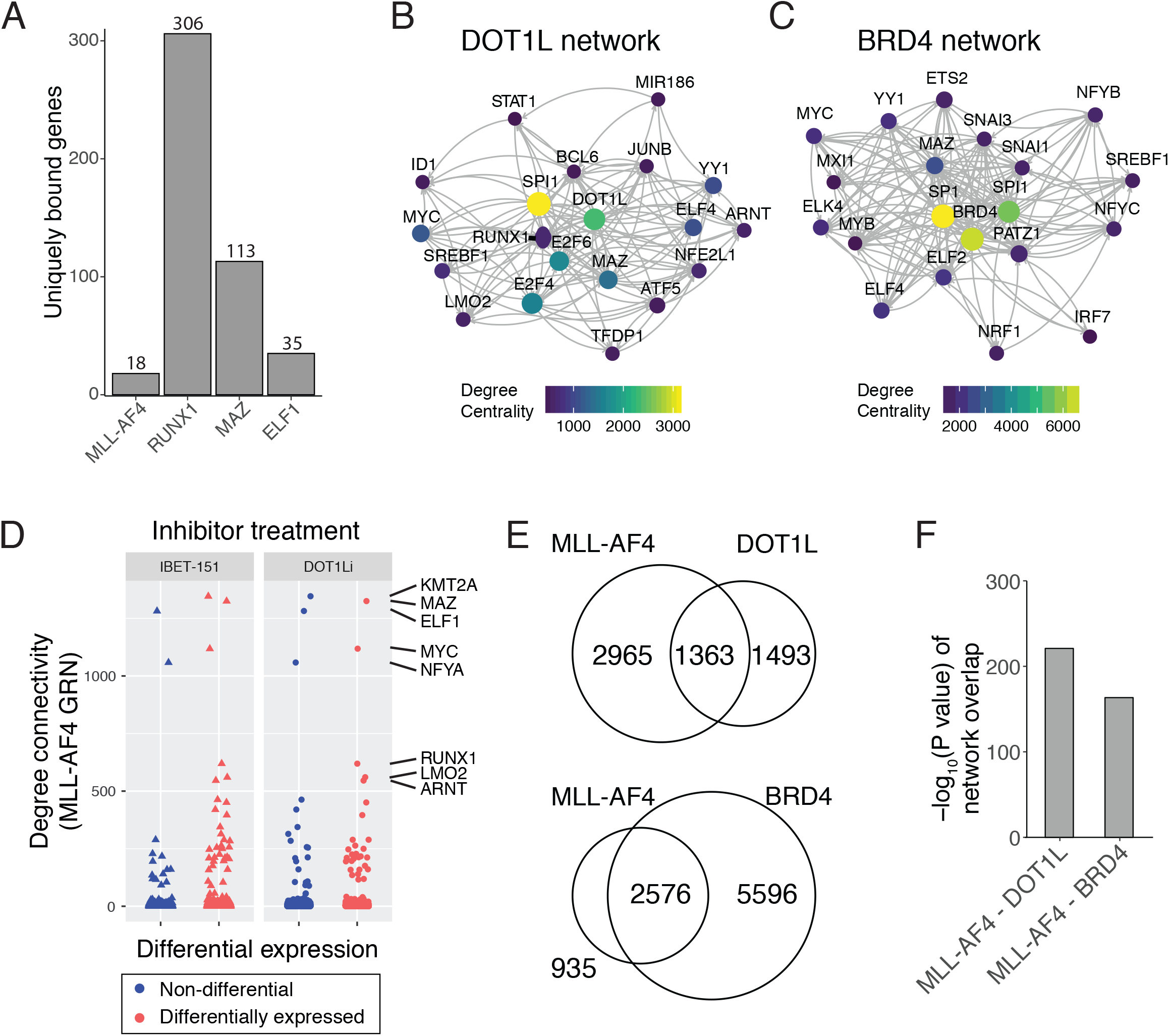
Developing GRN models for DOT1L and BRD4 (**A**) Quantification of the number of genes uniquely annotated in each ChIP-seq experiment in comparison with the other datasets. (**B-C**) Top 20 genes of the DOT1L (**B**) and BRD4 (**C**) GRN models by degree centrality. Lines (edges) indicate predicted interaction from protein (source) to gene locus (target), with arrowheads pointing towards downstream nodes. (**D**) DEGs by nascent RNA-seq upon 1.5 hours’ IBET or 7 days’ EPZ-5676 treatment, plotted against degree centrality in SEM MLL-AF4 GRN. (n=3). (**E**) Venn diagram overlaps between MLL-AF4, DOT1L and BRD4 GRNs. (**F**) Statistical significance of overlaps shown in (**E**). Significance calculated using a Fisher’s exact test and displayed as −log_10_ P value.

**Supplementary Figure 2.**
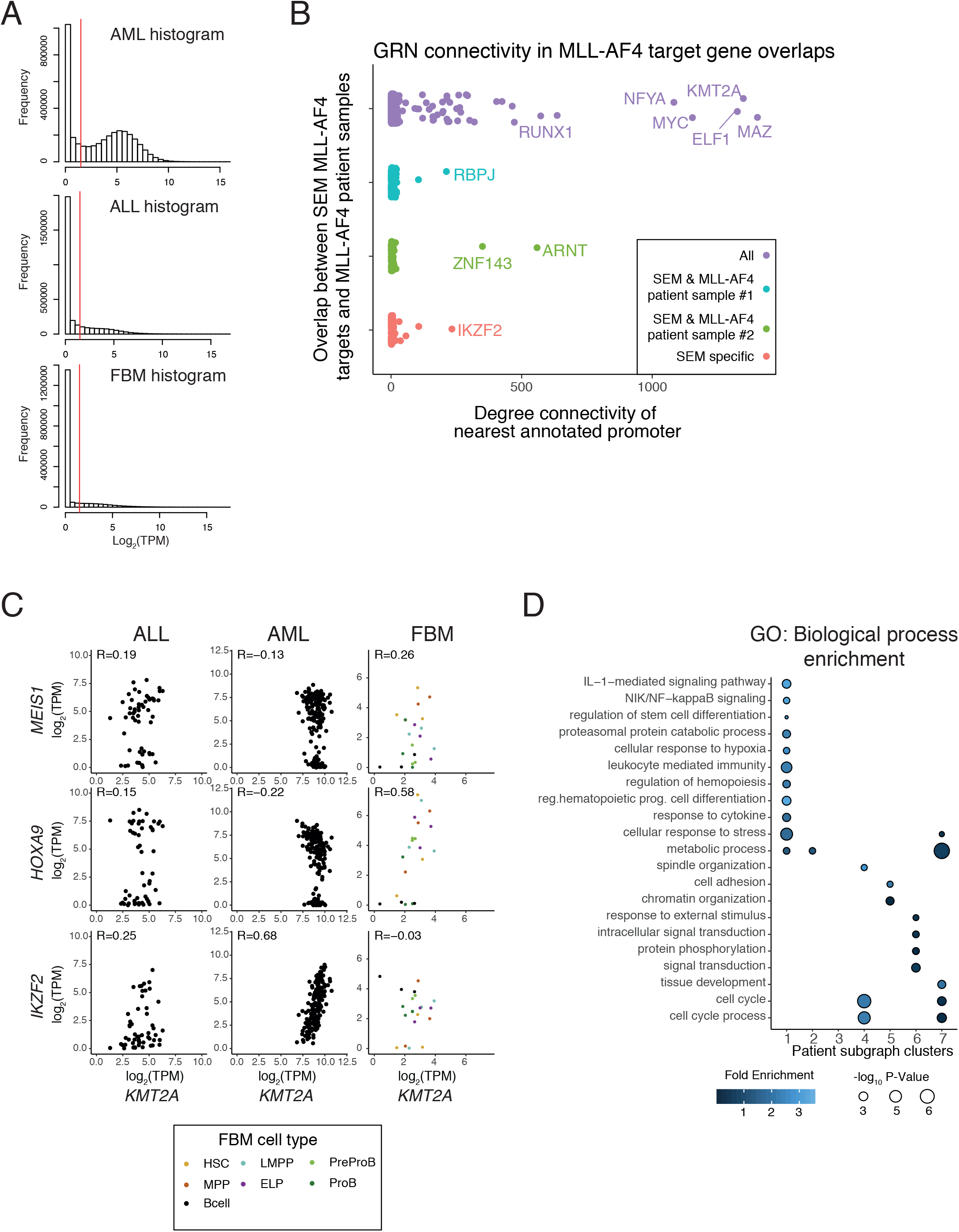
Patient sub-network analysis highlights core transcription factors. (**A**) Histograms of log2 TPM expression across AML, ALL and FBM datasets (67, 69, 70). Red line indicates 1.5 log2 TPM, the threshold used to define active expression. (**B**) Overlap groups of SEM MLL-AF4 target genes, as in Figure 1B, against degree connectivity within the SEM MLL-AF4 GRN. (**C**) Scatter plots of select genes, plotted from log2(TPM) expression data from the ALL, AML and FBM datasets. (**D**) GO Biological process enrichment for patient subgraph clusters. Size of points represents −log_10_ FDR of enrichment, while point color represents fold enrichment over expected number of genes.

**Supplementary Figure 3.**
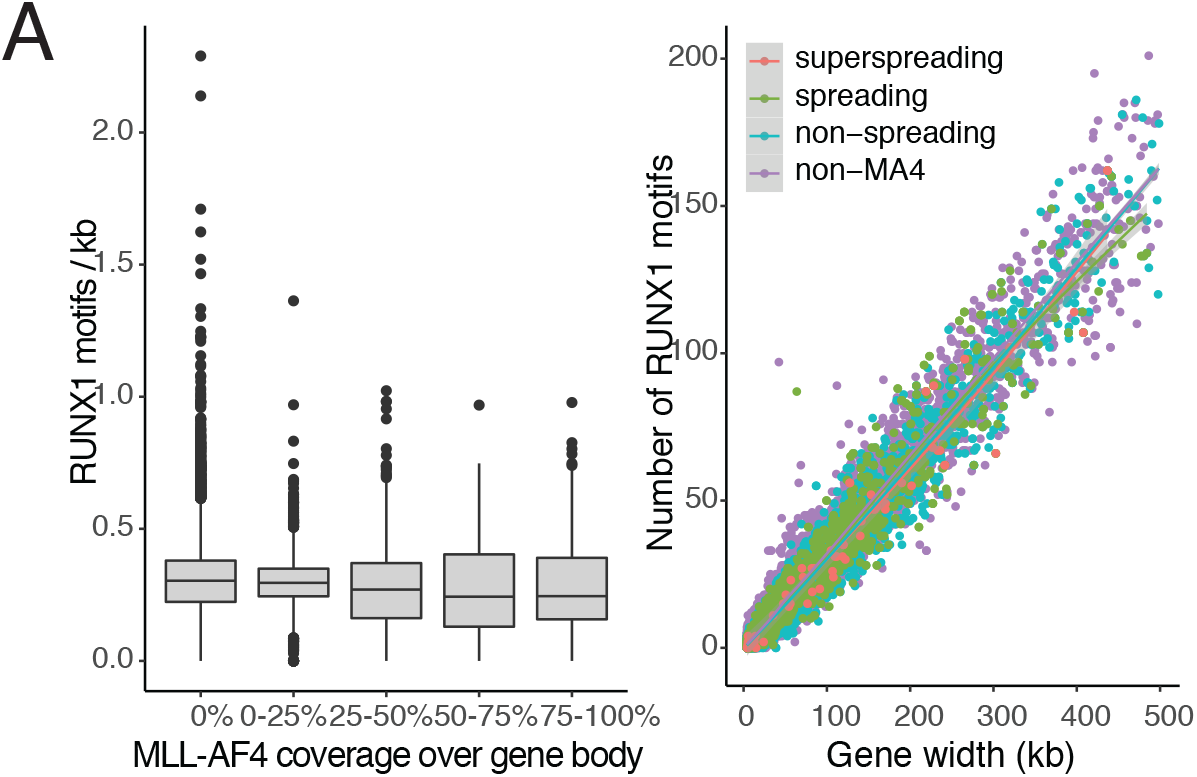
Canonical RUNX1 DNA binding motifs are not associated with MLL-AF4 binding profiles. (**A**) Left – Boxplots showing quantification of RUNX1 motifs/kb in a gene body by MLL-AF4 percent coverage of the gene body. Right – Scatter plot of number of RUNX1 motifs in a gene body against gene width (kb). Local regression lines (LOESS) added to describe the relationship between number of motifs and gene width, with grey ribbon representing confidence intervals. Local regression lines calculated for genes with no MLL-AF4, MLL-AF4 non-spreading, MLL-AF4 spreading greater than 5 kb, and MLL-AF4 spreading greater than 50 kb

**Supplementary Figure 4.**
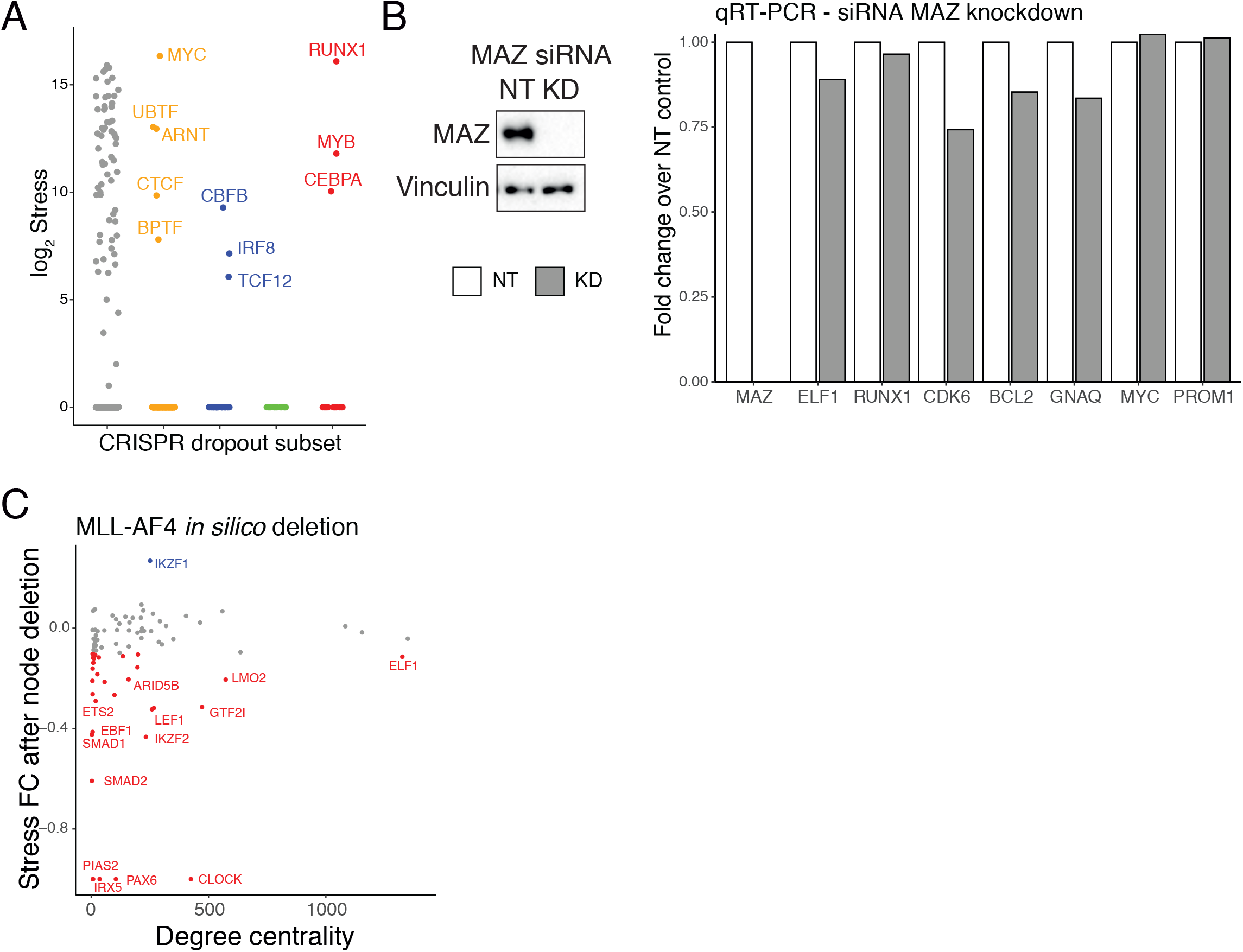
Comparison of MLL-AF4 GRN nodes with published CRISPR essentiality screens. (**A**) Association of Log_2_ stress centrality with CRISPR essentiality groups as outlined in Fig. 1A. G genes identified as essential in HT-29 (colon adenocarcinoma) or HT-1080 (Fibrosarcoma) were classified as non-specific essential. Genes not essential in HT-29 or HT-1080 but essential in MOLM-13 (MLL-AF9) or MV4-11 (MLL-AF4) were annotated as essential specifically for either MOLM-13 or MV4-11, or both cell lines. (**B**) Left - Western blot analysis in SEM cells probing MAZ protein after 96 hours’ siRNA knockdown (KD), or non-targeting (NT) siRNA treatment. Vinculin used as loading control. Right – qRT-PCR results showing expression of key MLL-AF4 targets after MAZ knockdown, relative to NT control. (n=1). Error bars represent standard error of the mean. (**C**) Stress fold change responses for individual transcription factors plotted against degree centrality for *in silico* deletion of MLL-AF4. Blue datapoints indicate positive stress fold change responses, while red indicate a negative response.

**Supplementary Figure 5.**
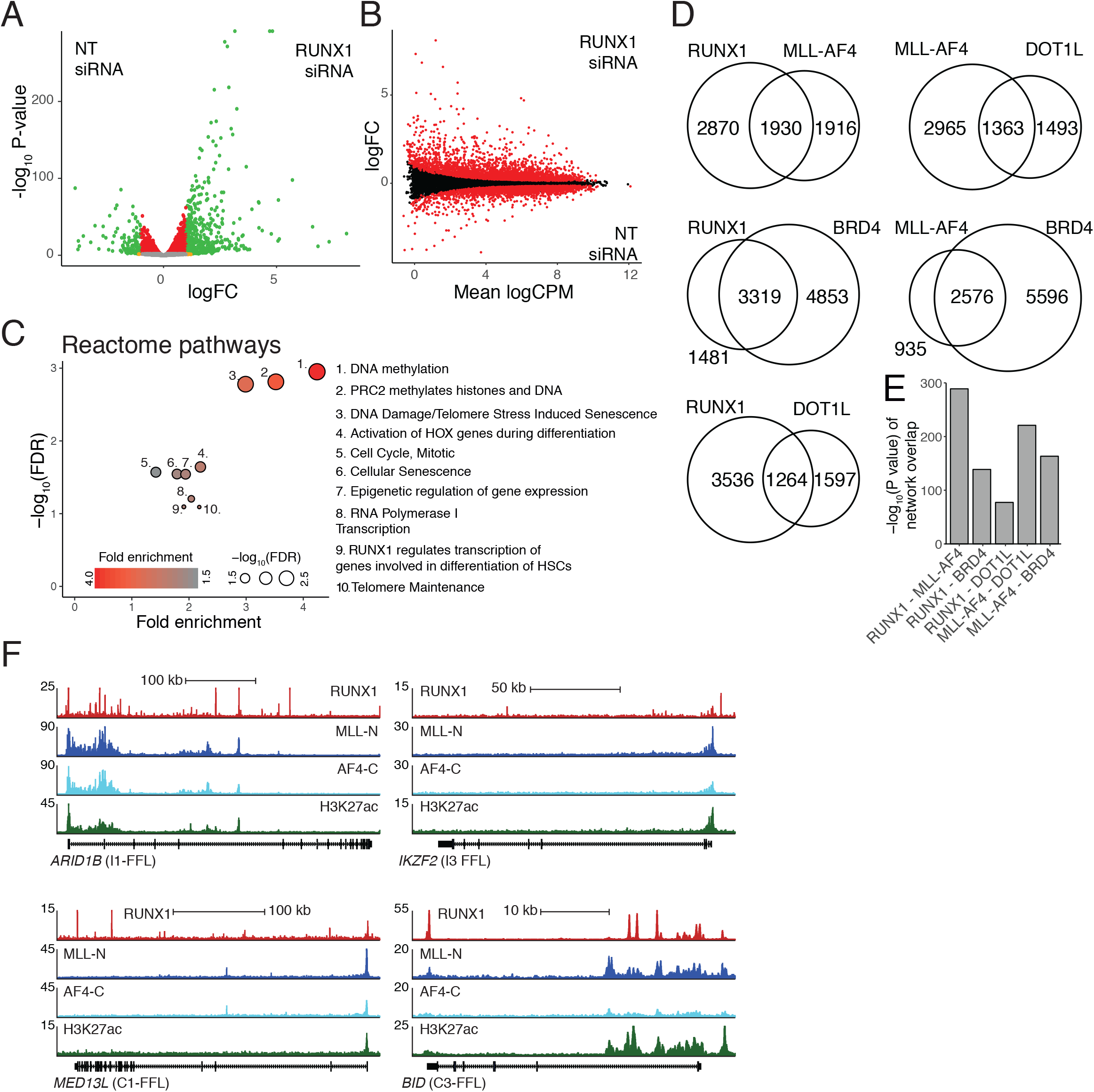
MLL-AF4 cooperates with RUNX1 in FFL and cascade circuits to regulate downstream targets. (**A**) Volcano plot showing the relationship between −log_10_ P value and |logFC| in RUNX1 siRNA knockdown nascent RNA-seq. Orange points represent genes |logFC| > 1 and FDR > 0.05; red represents |logFC| < 1 and FDR < 0.05; green represents |logFC| > 1 and FDR < 0.05. (**B**) MA plot showing the relationship between logFC and mean log_2_ CPM in RUNX1 siRNA knockdown nascent RNA-seq. DEGs are highlighted in red. (**C**) Pathway enrichment (Reactome) for overlap between MLL-AF4 and RUNX1 KD DEGs (Figure 5B). Size of points represents −log_10_ FDR of enrichment, while point color represents fold enrichment over expected number of genes. (**D**) Venn diagram overlaps between RUNX1, MLL-AF4, DOT1L and BRD4 GRNs. (**E**) Statistical significance of overlaps shown in (**D**). Significance calculated using a Fisher’s exact test and displayed as −log_10_ P value. (**F**) ChIP-seq tracks generated in hg19 for MLL-N, AF4-C, RUNX1 and H3K27ac. ChIP-seq data normalized to 1×10^7^ reads. Displayed loci are for MLL-AF4 FFL circuit targets.

**Supplementary Figure 6.**
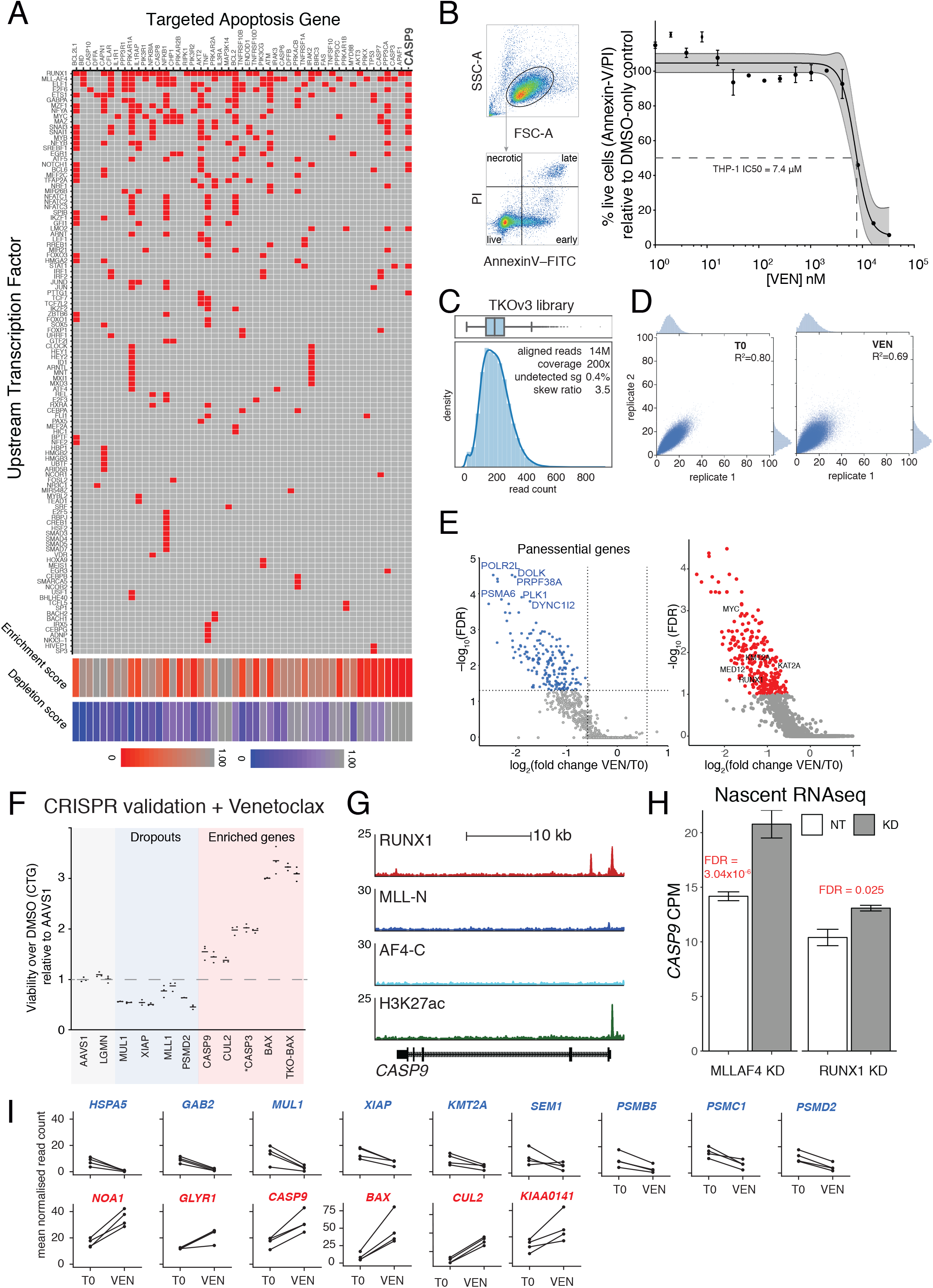
CRISPR screen in combination with venetoclax treatment to test GRN predicted circuits. (**A**) Interaction matrix showing TF regulators of apoptosis pathway (rows) and components of the apoptosis pathway (columns). Select subset of TFs and pathway components shown. Red squares indicate a predicted regulatory interaction between the TF and apoptosis component. (**B**) Left – Flow cytometry gating for Annexin-V/PI staining of THP-1 cells to assay viability with venetoclax treatment. Right – THP-1 cells were cultured for 48 hours with different concentrations of venetoclax, and assayed for viability. PI: propidium iodide; early/late: early/late apoptotic. Dots show mean; error bars show SD. Lines show unconstrained four parameter sigmoidal least-squares regression best fit; shaded region shows 99% confidence interval. For all curves, R2 > 0.99. (n=5). (**C**) Sequencing validation of amplified sgRNA pool showing sgRNA distribution as a histogram and boxplot. (**D**) Scatter plot of sgRNA counts for biological replicates of T0 and +VEN samples. Correlation between replicates shown as R2, and histograms distributions shown along the axis. (**E**) Analysis of CRISPR screen using MAGeCK, plotting −log_10_ FDR against log_2_ gRNA fold change. Left – MAGeCK results, including only pan-essential genes. Right – MAGeCK results for depleted sgRNA, filtered for MLL-AF4 GRN nodes. (**F**) Functional validation of CRISPR screen hits by individual gene knockouts using sgRNA from the Brunello library. sgRNA were cloned into lentiCRISPRv2 constructs and transduced into THP-1 cells, followed by 48h treatment with DMSO or 20 μM Venetoclax. CellTiter-Glo (CTG) was used to assay viability, and is normalized as Venetoclax/DMSO viability, relative to AAVS1 silent control (71). Genes marked * are FDR > 0.1 in the CRISPR screen. Dots show biological replicates (n=3); bars show mean. (**G**) Expression of *CASP9* from nascent RNA-seq data following 96 hours MLL-AF4 knockdown. Expression normalized as CPM. (**H**) ChIP-seq tracks generated in hg19 for MLL-N, AF4-C, RUNX1 and H3K27ac at the *CASP9* locus. ChIP-seq data normalized to 1×10^7^ reads. (**I**) Normalized mean sgRNA counts for key depleted and enriched genes at T0 and T18+Venetoclax (n = 2). Dropout hits are highlighted in blue, enriched in red.

**Supplementary Table S1.**
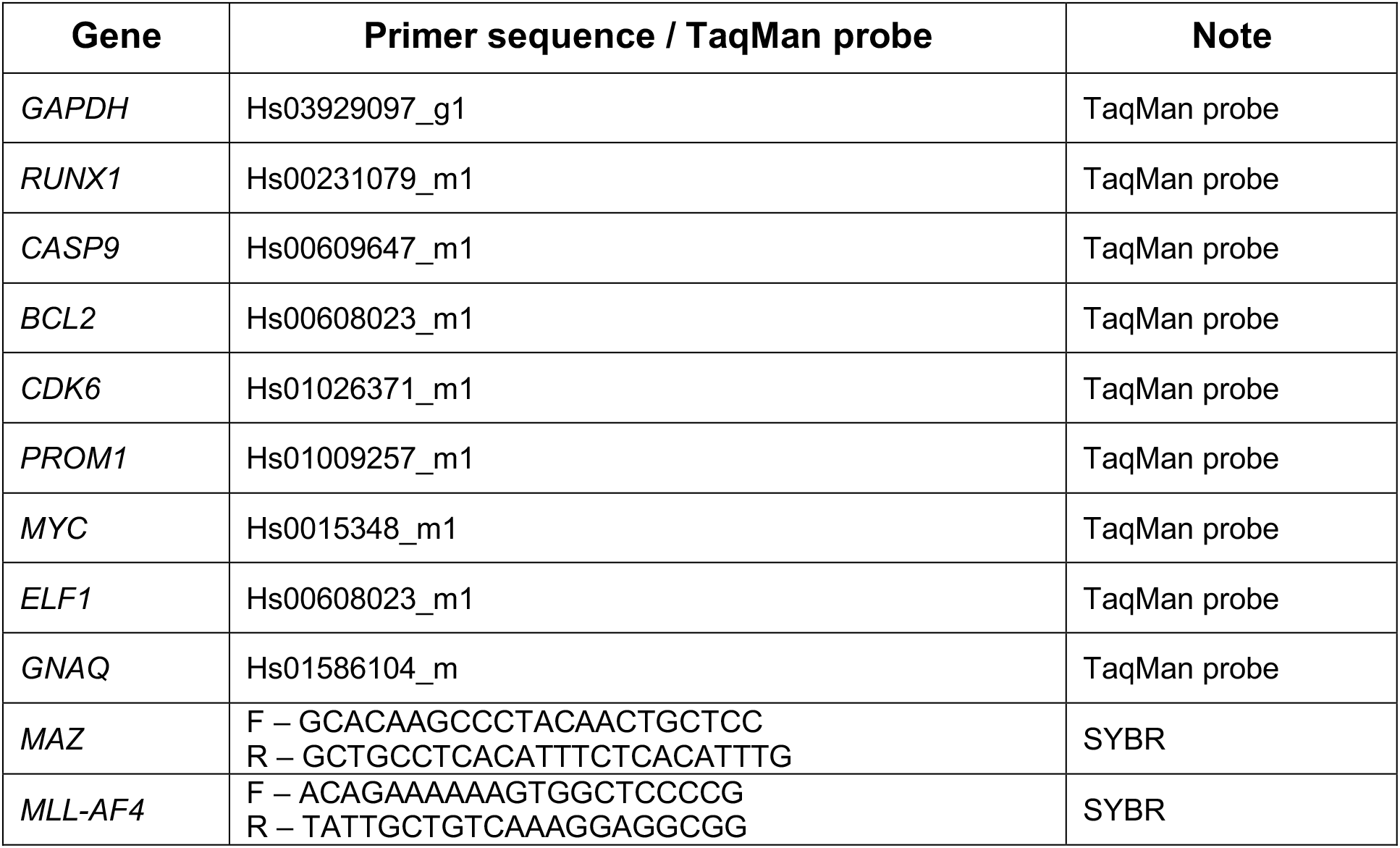
List of qRT-PCR primers

**Supplementary Table S2.** GEO accession numbers for previously published sequencing experiments

| **Experiment** | **Cell line/tissue** | **Antibody** | **Treatment** | **Accession Number** |
| --- | --- | --- | --- | --- |
| ChIP-seq | SEM | MLL-N |  | GSE74812 |
| ChIP-seq | SEM | AF4-C |  | GSE74812 |
| ChIP-seq | SEM | RUNX1 |  | GSE42075 |
| ChIP-seq | SEM | ELF1 |  | GSE117865 |
| ChIP-seq | SEM | H3K79me3 |  | GSE117865 |
| ChIP-seq | SEM | BRD4 |  | GSE83671 |
| ChIP-seq | SEM | H3K27ac |  | GSE74812 |
| ATAC-seq | SEM |  |  | GSE74812 |
| ChIP-seq | Primograft | MLL-N |  | GSE83671 |
| ChIP-seq | Primograft | AF4-C |  | GSE83671 |
| Nascent RNA-seq | SEM |  | MLL-AF4 siRNA | GSE85988 |
| Nascent RNA-seq | SEM |  | EPZ-5676 | GSE83671 |
| Nascent RNA-seq | SEM |  | IBET | (67) |

## Description of Additional Supplementary Files

File Name: Supplementary Data 1

Description: MLL-AF4 and RUNX1 annotated edge and node tables.

File Name: Supplementary Data 2

Description: Patient sub-network node clusters

File Name: Supplementary Data 3

Description: Nascent RNA-seq differential genes after RUNX1 siRNA knockdown and PANTHER pathway enrichment of DEGs in common between MLL-AF4 and RUNX1 siRNA treatment.

File Name: Supplementary Data 4

Description: MLL-AF4 – RUNX1 regulatory FFL and cascade circuits.

File Name: Supplementary Data 5

Description: Results of CRISPR screen in combination with venetoclax treatment in THP-1 cells.

